# CREB REGULATES FOXP3^+^ST-2^+^ TREGS WITH ENHANCED IL-10 PRODUCTION

**DOI:** 10.1101/2024.06.29.601312

**Authors:** Sudheendra Hebbar Subramanyam, Judit Hriczko, Saskia Schulz, Thomas Look, Tannaz Goodarzi, Tim Clarner, Miriam Scheld, Markus Kipp, Eva Verjans, Svenja Böll, Christopher Neullens, Ivan Costa, Zhijian Li, Lin Gan, Bernd Denecke, Angela Schippers, Stefan Floess, Jochen Huehn, Edgar Schmitt, Tobias Bopp, Hermann Wasmuth, Ron Winograd, Rudi Beyaert, Bart Lambrecht, Martin Zenke, Norbert Wagner, Kim Ohl, Klaus Tenbrock

**Affiliations:** Dept. of Pediatrics, RWTH Aachen University; Institute for Biomedical Engineering, Department of Cell Biology, RWTH Aachen, University Hospital, Aachen, Germany; Helmholtz Institute for Biomedical Engineering, RWTH Aachen University, Aachen, Germany; Institute of Neuroanatomy and JARA-BRAIN, Faculty of Medicine, RWTH Aachen University; Institute of Anatomy, Medical University of Rostock, Rostock, Germany; Instutute for Computational Genomics, IZKF, RWTH Aachen University; Genomics Facility, Interdisciplinary Center for Clinical Research Aachen (IZKF Aachen), RWTH Aachen University, Aachen, Germany; Department of Experimental Immunology, Helmholtz Centre for Infection Research, Braunschweig, Germany; Institute for Immunology, University Medical Center, Johannes Gutenberg University Mainz, Mainz, Germany; Luiusenhospital Aachen, Dept. of Medicine, Aachen, Germany; VIB Center for Inflammation Research, Ghent, Belgium; Department of Biomedical Molecular Biology, Ghent University, Ghent, Belgium; Department of Respiratory Medicine, Ghent University Hospital, Ghent, Belgium; Division of Pediatric Rheumatology, Department of Pediatrics, Inselspital, Bern University Hospital, University of Bern, Switzerland

**Author notes:** Corresponding author: Klaus Tenbrock, Dept. of Pediatrics, RWTH Aachen University, Pauwelsstraße 30, 52074 Aachen, 0049-2418089140. Bernd Denecke unfortunately passed away during the project. equal contribution (Kim Ohl, Klaus Tenbrock). The authors have declared that no conflict of interest exists.

## Abstract

Regulatory T cells (T_regs_) are gatekeepers of immune homeostasis and characterized by expression of Foxp3, which maintains T_reg_ identity. Here we demonstrate that in mice with a Foxp3-specific knockout of CREB, enhanced numbers of T_regs_ are found *in vivo* in spleen, lung and colon. These T_regs_ display a reduced Foxp3 expression, but enhanced expression of the IL-33 receptor (ST-2), IL-10, IL-13, and CREM. CREB deficient T_regs_ were highly suppressive *in vitro* and prevented disease activity in CD4 T cell mediated transfer colitis in an IL-10 dependent way. Mechanistically CREB fulfils dual roles in T_regs_. First it downregulates Foxp3 expression, however in cooperation with CREM, CREB expression in T_regs_ alters chromatin accessibility to the ST-2 region and thereby influences T cell specific immune responses mediated by IL-10.

Brief summary: Mice with a Foxp3-specific knockout of CREB display enhanced expression of IL-13, IL-10, ST-2 and CREM, which prevents gut inflammation

**GRAPHICAL ABSTRACT:** 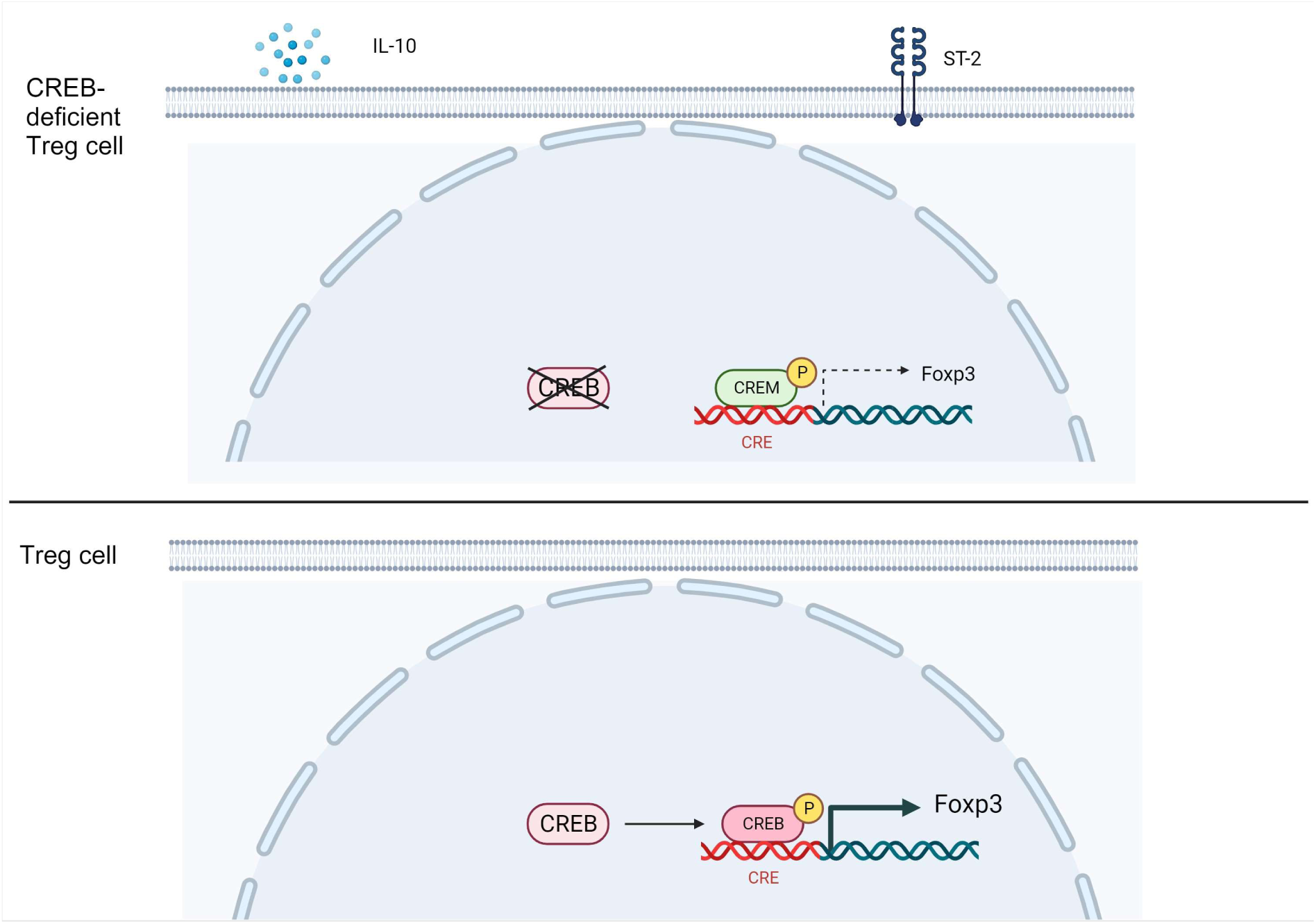

Created by Biorender

## 1 INTRODUCTION

Regulatory T cells (T_regs_) are important for the maintenance of immune homeostasis and characterized by expression of Foxp3. Loss of Foxp3 results in a devastating autoimmune syndrome in mice (scurfy disease)(Wildin, Ramsdell et al. 2001) and humans (IPEX, polyglandular autoimmunopathy) (Bennett, Christie et al. 2001). Two sources of Foxp3^+^ cells have been described, one of which is located to the thymus, “thymus-derived T_regs_” (tT_regs_), and the second to the periphery, where conventional CD4^+^ T cells can convert into Foxp3^+^ “inducible T_regs_” (iT_regs_) after T cell stimulation in the presence of TGF-β (Zhou, Bailey-Bucktrout et al. 2009). tT_reg_-cell as well as iT_reg_ cell generation largely depend on the presence of IL-2 which is not only required for homeostatic maintenance but also for the thymic development of T_regs_ (Setoguchi, Hori et al. 2005, Zorn, Nelson et al. 2006, Liao, Lin et al. 2011, Ohl and Tenbrock 2015).

ST-2 also known as IL1RL1 belongs to Toll-like receptor/IL-1 receptor superfamily. ST-2 through its ligand IL-33 plays a pivotal role in MyD88/ NFκB signaling. The interaction of ST-2 with its ligand IL-33 induces Foxp3 and GATA3 expression in T_regs_ in particular at mucosal sites (Schiering, Krausgruber et al. 2014). ST-2 mainly exists in two isoforms; soluble ST-2 (sST-2) and membrane bound ST-2. sST-2 functions as a decoy receptor of IL-33 and inhibits NFκB signaling, while membrane bound ST-2 promotes NFκB signaling (Kakkar and Lee 2008). It has been shown that increased ST-2 levels were observed in diseases like inflammatory bowel disease (IBD), Asthma, colon cancers, and Graft versus Host disease (Griesenauer and Paczesny 2017). In many non-lymphoid tissues, T_regs_ show enriched expression of the ST-2 and most of these are iT_regs_. The regulation of the ST-2^+^T_regs_ is dependent on the transcription factor Basic leucine zipper transcription factor ATF-like (BATF) (Delacher, Imbusch et al. 2017, Delacher, Imbusch et al. 2020), while other factors which have not been defined yet. BATF belongs to the cAMP responsive element binding protein/activating transcription factor (CREB/ATF) family of transcription factors, is able to dimerize with Jun and has been shown to compete with CREB for binding to CRE/AP1 sites (Dorsey, Tae et al. 1995, Manna and Stocco 2007).

The accessibility of the *Foxp3* gene promoter, as well as other cis-regulatory elements in the *Foxp3* locus are under tight epigenetic control, especially by histone acetylation, methylation and cytosine-phospho-guanosine (CpG-)DNA methylation (Lal and Bromberg 2009). The *Foxp3* gene contains a highly conserved CpG-rich region in the first intron (+4201 to +4500) which appears to be critical in *Foxp3* gene regulation. CpG residues in this region are completely methylated (thus resulting in a “silenced” gene locus) in non-T_regs_ (CD4^+^CD25^-^) and fully de-methylated (“open, accessible” gene locus) in tT_regs_ (CD4^+^CD25^+^) in mice and humans, for which this region was termed T_reg_-specific de-methylated region (TSDR) (Baron, Floess et al. 2007, Floess, Freyer et al. 2007, Kim and Leonard 2007). The TSDR possesses transcriptional enhancer activities and transcription factors c-Rel, Ets-1 and those of the CREB/ATF family have been identified to bind and trans-activate the de-methylated TSDR (Floess, Freyer et al. 2007, Kim and Leonard 2007, Polansky, Schreiber et al. 2010). Thus, CREB was identified previously as a positive regulator of Foxp3 expression *in vitro*.

Through yet-to-be-defined mechanisms, T_regs_ loose Foxp3 expression and become pro-inflammatory effector cells under certain conditions (Tsuji, Komatsu et al. 2009, Zhou, Bailey-Bucktrout et al. 2009). In detail the loss of Foxp3 expression results in T_h1_, T_h17_ and T follicular helper (T_fh_) conversion of T_regs_, while a reduction of Foxp3 expression leads to a T_h2_ conversion of tT_regs_ (Wang, Souabni et al. 2010). Therefore, additional molecular events might complement Foxp3 function in the generation of T_regs_ and the maintenance of their function and phenotype. DNA methylation is one of the important epigenetic regulations to control transcription. As mentioned above, methylation patterns of Foxp3 promoter and enhancer regions have been intensively investigated in the past years. In addition, a paper by the group of Feuerer showed tissue specific regulation of DNA methylation in T_regs_ also regarding other loci including gene sites that are associated with the T_h2_ subset (Delacher, Imbusch et al. 2017).

Our understanding of molecular mechanism of suppression is still limited. T_regs_ control immune activation by acting directly or indirectly on effector CD4 and CD8 T cells and antigen-presenting cells. Cyclic AMP (cAMP) is long known as a potent suppressor of T cell activation and function. T_regs_ generate and accumulate high levels of cAMP, which can be transferred into target cells as one mechanism of suppression (Bopp, Becker et al. 2007, Bodor, Bopp et al. 2012). Indirect mechanisms include the secretion of cytokines, such as IL-10, IL-35 and TGF-ß. IL-10 plays an important role in suppressing CD4^+^ effector T-cell function. Additionally, IL-10 prevents activation of antigen-presenting cells (APCs), such as dendritic cells and macrophages, by downregulation of costimulatory molecules and production of inflammatory cytokines within these cells (Okamoto, Fujio et al. 2011). Selective ablation of IL-10 in Foxp3^+^ T_regs_ revealed that IL-10 production by T_regs_ is essential for keeping the immune response in check at environmental interfaces such as colon and lungs (Rubtsov, Rasmussen et al. 2008). IL-10 also acts in an autocrine manner by maintaining FoxP3 expression and the suppressive capacity of T_regs_ (Murai, Turovskaya et al. 2009, Unutmaz and Pulendran 2009). Interestingly, IL-10 is responsive to cAMP stimulation at least in THP1 cells and macrophages and the IL-10 promoter contains four putative cAMP response element sites (CRE, TGACGTCA) sites, which are conserved between men and mice (Brenner, Prosch et al. 2003). Following TLR activation, pCREB (phosphorylated CREB) is recruited to the IL-10 promoter and enhances IL-10 transcription in macrophages (Sanin, Prendergast et al. 2015).

CREB is a phosphorylation-dependent transcription factor that is involved in several cellular processes, amongst others proliferation and cell survival. CREB and CREM belong to the CREB/activating transcription factor (ATF) family. Hallmark of which is a conserved basic region/leucine zipper (bZIP) domain, which binds to an 8-base pair palindromic DNA sequence, called CRE site (Foulkes, Borrelli et al. 1991, Sassone-Corsi 1995, De Cesare and Sassone-Corsi 2000). The transactivation domain is required for the recruitment of CREB binding protein (CBP/p300), which has histone-acetylating (HAT) activities. CREM has many alternative splice variants. ICER is a splice variant of CREM that lacks transcriptional activation domains and thereby functions as a repressor of cAMP-induced CRE-mediated transcription. CREMα also lacks both Q-domains. The Q2 domain is needed to activate HAT activities of p300 and CBP and CREMα is therefore unable to promote HAT activities of p300 and CBP (Asahara, Santoso et al. 2001). ICER also has an alternative transcription initiation site and is induced by a unique alternative promoter (Foulkes, Borrelli et al. 1991). ICER has been described as central to Th17 cell differentiation in autoimmunity (Yoshida, Comte et al. 2016). On the other hand, Bodor et al. showed that ICER is important for cAMP-driven Treg-cell suppression mechanism in activated T cells (Bodor, Bopp et al. 2012). Expression of CREMα, a repressor isoform of CREM, is increased in CD4^+^ T cells from SLE patients, and forced expression of CREMα in human T cells enhances IL-17A expression (Rauen, Hedrich et al. 2011). Moreover, mice overexpressing CREMα in T cells display increased IL-17 production and severe skin inflammation, as well as mild lupus-like disease (Lippe, Ohl et al. 2012). In addition, we have shown that CREMα actively recruits a histone deacetylase (HDAC1) to promote chromatin condensation (Tenbrock, Juang et al. 2006). As a third mechanism CREMα recruits the DNA methyltransferase DNMT3a, which promotes methylation of DNA and thereby represses the binding of transcription factors resulting in closure of chromatin (Hedrich, Rauen et al. 2011). CREMα itself can furthermore bind to the aforementioned CRE within the *Il2* promoter (Powell, Lerner et al. 1999), (Lippe, Ohl et al. 2012, Ohl, Wiener et al. 2015). The group of George Tsokos demonstrated that SLE T cells have a decreased ability to produce IL-2 in response to stimulation with antigens (Via, Tsokos et al. 1993), which is related to enhanced T cellular expression of CREMα in these patients and an aberrant CREM/CREB ratio (Solomou, Juang et al. 2001). CREB binds to a CRE site within the proximal *IL2* promoter resulting in enforced *IL2* transcription. Mice expressing a dominant negative (inactive) CREB isoform show a markedly decreased production of IL-2. Additionally, CREB^-/-^ mice die prenatally while they display a severely impaired T cell development (Barton, Muthusamy et al. 1996).

Previous data from Wang et al., group suggested that CREB negatively regulated the survival of iT_regs_ and is particularly important in the generation of CD4+Th17 cells. Using CD4^CRE^ CREB^fl/fl^ mice they showed that these cells prevented colitis in a Rag2^-/-^ model, however their model has the disadvantage that CD4 cells of these mice are expressing lower amounts of IL-2 that could hamper iT_reg_ induction. Our analysis now expands their findings and determines an important tole for CREB in the generation of ST2+T_regs_. Using Foxp3^CRE^CREB^fl/fl^ mice we herein confirm the colitis phenotype but can show that i) CREB deficiency in T_regs_ induces expression of ST2+T_regs_, ii) the phenotype that prevents colitis in Rag2^-/-^ mice is primarily dependent on abundance of IL-10, iii) and that this phenotype can be reversed by knockout of CREM that lowers the expression of ST2.

## 2 MATERIAL AND METHODS

### 2.1 Mice strains

Experiments were performed with age matched *CREB^fl/fl^*and *Foxp3^cre^CREB^fl/fl^* mice (all C57BL/6). *Foxp3^cre^CREB^fl/fl^*mice were generated by crossing CREB-flox mice (Okawa, Motohashi et al. 2006) with C57BL/6 Foxp3-IRES Cre mice (provided by T. Bopp, University of Mainz). *Foxp3^cre-^CREB^fl/fl^* mice were used as controls (denoted as *CREB^fl/fl^*). Foxp3^Cre^ROSA^RFP^ were provided by T. Bopp University of Mainz and crossed with our *CREB^fl/fl^*mice. Rag2^-/-^ (C57BL/6) mice were provided by A. Schippers (Uniklinik RWTH Aachen). All mice were bred in our animal facility and kept under standardized conditions.

### 2.2 Transfer colitis

To induce transfer colitis, Rag2^-/-^ mice were adoptively transferred with 2 x 10^6^ CD4^+^ CD25^-^ T cells. Animals were sacrificed as soon as a significant loss of weight was measurable. Spleen and mLNs were harvested for further analysis. One part of the colon was fixed in formalin for histological scoring and the other part was fixed in RNAlater (Qiagen, Germany) for subsequent mRNA analysis. For soluble ST2 (sST2) treatment, mice received 100 µg mouse sST2 (produced by the VIB Protein Core facility and provided by R. Beyaert, VIB-UGent Center for Inflammation Research), three times/week by intraperitoneal means (i.p). Control group received PBS i.p. For IL-10R treatment, mice received 500 µg anti IL-10R antibody (1B1mAB) (kindly provided by Prof. Edgar Schmitt, University of Mainz) or IgG control AK (Rat IgG kappa) i.p. once a week during the first three weeks.

### 2.3 Histological scoring

4 µm paraffin sections from the fixed colon were cut serially, mounted onto glass slides, and deparaffinized. The colon sections were stained with hematoxylin and eosin by the Core Facility (IZKF) of the RWTH Aachen University. Blinded histological scoring was performed using a standard microscope, based on JLS method as described previously (Bleich, Mahler et al. 2004, Pils, Bleich et al. 2011). Each colon section was scored for the four general criteria: severity, degree of hyperplasia, degree of ulceration, if present, and percentage of area involved. A subjective range of 1-3 (1 = mild, 2 = moderate, 3 = severe) was used for the first three categories. Severity: Focally small or widely separated multifocal areas of inflammation limited to the lamina propria were graded as mild lesions (1). Multifocal or locally extensive areas of inflammation extending to the submucosa were graded as moderate lesions (2). If the inflammation extended to all layers of the intestinal wall or the entire intestinal epithelium was destroyed, lesions were graded as severe (3). Hyperplasia: Mild hyperplasia consisted of morphologically normal lining epithelium that was at least twice as thick (length of crypts) as adjacent or control mucosa. Moderate hyperplasia was characterized by the lining epithelium being two-or three-times normal thickness, cells were hyperchromatic, numbers of goblet cells were decreased, and scattered individual crypts developed an arborizing pattern. Severe hyperplastic regions exhibited markedly thickened epithelium (four or more times normal thickness), marked hyperchromasia of cells, few to no goblet cells, a high mitotic index of cells within the crypts, and numerous crypts with arborizing pattern. Ulceration was graded as: 0 = no ulcer, 1 = 1-2 ulcers (involving up to a total of 20 crypts), 2 = 1-4 ulcers (involving a total of 20-40 crypts), and 3 = any ulcers exceeding the former in size. A 10% scale was used to estimate the area involved in the inflammatory process. 0 = 0%, 1 = 10%-30%, 2 = 40%-70%, 3 = >70%.

### 2.4 Cell isolation

Single cell suspensions were isolated from spleens and LNs (pLN and mLNs) using cell strainers and erythrocytes were lysed with lysis buffer. To obtain immune cells from lung, tissue was excised into small pieces and was digested with 0.1% collagenase in DPBS for 1hr. The digested fraction was passed through a 40μm nylon strainer and rinsed with DPBS. The flow through was centrifuged and the cell pellet was lysed using RBC lysis buffer. The cell pellet was used for the Flow cytometric analysis. Colon was retrieved from the mice and rinsed with a cannula with PBS+0.5%BSA. Afterwards colon was excised into small pieces and placed in a digestive medium containing RPMI (10ml), Collagenase V (0.85mg/ml), Collagenase D (1.25mg/ml), Dispase (1mg/ml) and DNase (30μg/ml) and incubated for 45 minutes at 37°C with gentle shaking. The digested fraction was passed through a 40um nylon strainer and the flow through was collected and centrifuged to obtain a pellet. Percoll gradient centrifugation was performed by using 35% percoll and centrifugation was performed. The cell pellet obtained was lysed using RBC lysis buffer and is used for further analysis. Liver was retrieved from the mice. Gall bladder was removed and liver was excised into small pieces and was placed in a digestive medium containing RPMI without FCS (3ml), 1.25mg/ml Collagenase D and 30μg/ml DNase. The fraction was incubated for 45 minutes and digestion was stopped by adding PBS+0.5% BSA+2mM EDTA and homogenized using a syringe. The digested fraction was passed through a 100μm nylon strainer and the flow through was centrifuged. The pellet obtained was used for the percoll gradient centrifugation by using 35% percoll. The cell pellet obtained after the percoll gradient centrifugation was lysed using RBC lysis buffer and is used for flow cytometric analysis.

### 2.5 T cell differentiation assays

MACS isolated CD4^+^CD25^-^ T cells (2×10^6^ per ml) were incubated with plate-coated anti-CD3 (10µg/ml, eBioscience) and soluble anti-CD28 (1µg/ml) (eBioscience). T_H_0 cells were left without exogenous cytokines, 5 ng/ml TGF-β was added to induce T_reg_ differentiation.

### 2.6 Flow cytometry

For surface staining, single cell suspensions were stained with anti-CD4, anti-CD3, anti-CD8, anti-B220, anti-CD25, CD45, anti-GL-7, anti-ICOS, anti-ST2, anti-Nrp1, anti-PD-1 (all from eBioscience, Germany). For intracellular staining of CREB, Foxp3, CTLA-4, Helios, GATA3, T-bet and RORγt, cells were fixed and permeabilized with a FOXP3 staining buffer set (Thermofischer, eBioscience, Germany) following the manufacturer’s instructions and stained with respective antibodies (eBioscience, Germany) for 30 min. Intracellular cytokines were stained with anti-IFN-γ-Alexa 647 (eBioscience) and anti-IL-17-Alexa 488 (BD), and IL-10-APC (ebiosciences) after PMA (30nM) and Ionomycin (1.5µM) (both Sigma-Aldrich, USA) re-stimulation in the presence of GolgiPlug/GolgiStop (BD Biosciences). A daily calibrated FACS-Canto II flow cytometer (Becton Dickinson, MountainView, CA) was used to perform phenotypic analysis. Lymphocytes were gated by forward (FSC), side scatter (SSC), and CD3, CD4, B220 expression. A figure exemplifying the gating strategies is provided in the Supplementary Information (**Supp. Fig. 5**). For data analysis FCS-Express 4.0 Research Edition (DeNovo software Glendale, CA, USA) and FlowJo version 10 were used.

### 2.7 Luciferase Assay

RLM cells carrying a stable pGL3-TSDR-FoxPro luciferase plasmid were kindly provided by Jochen Huehn (HZI Braunschweig). Cells were transfected with pcDNA-CREB plasmid or pcDNA. Cells were left to rest over night before luciferase activity was measured by using the Dual-Glo Luciferase Assay System (Promega, USA).

### 2.8 RNA isolation and real-time PCR

Total RNA from isolated T cells and colon tissue was isolated using the RNeasy Mini Kit (Qiagen, Germany). cDNA was then generated from 200 ng total RNA using the RevertAid H Minus First Strand cDNA Synthesis Kit (Thermo Fisher Scientific, USA) according to the manufactureŕs instructions. RT-PCR was performed using the SYBR Green PCR kit (Eurogentec, Germany) and data were acquired with the ABI prism 7300 RT-PCR system (Applied Biosystems / Life Technologies, Germany). Each measurement was set up in duplicate. After normalization to the endogenous reference control gene ß-actin for mice, the relative expression was calculated.

### 2.9 RNA extraction and microarray for gene expression analysis

CD4^+^CD25^+^Nrp1^+^ T_regs_ were sorted by FACS. Genome wide transcriptome analyses of Foxp*^cre^CREB^fl/fl^* and *CREB^fl/fl^* T_regs_ were performed in independent triplicates using Gene Chip® Mouse Gene 2.0 arrays (Affymetrix, Santa Clara, CA, USA). Total RNA extraction was carried out using the RNeasy Micro Kit (Qiagen, Germany) according to manufacturer’s protocol and then quantified (Nanodrop). RNA quality was assessed using the RNA 6000 Nano Assay with the 2100 Bioanalyzer (Agilent, Santa Clara, CA, USA). Samples for the Gene 2.0 arrays were prepared and hybridized to the arrays according to the Affymetrix WT Plus Kit manual. Briefly, for each sample, 100 ng of total RNA was reversed transcribed into cDNA using a random hexamer oligonucleotide tagged with a T7 promoter sequence. After second strand synthesis, double strand cDNA was used as a template for amplification with T7 RNA polymerase to obtain antisense cRNA. Random hexamers and dNTPs spiked out with dUTP were then used to reverse transcribe the cRNA into single stranded sense strand cDNA. The cDNA was then fragmented with uracil DNA glycosylase and apurinic/apyrimidic endonuclease 1. Fragment size was checked using the 2100 Bioanalyzer and ranged from 50 to 200 bp. Fragmented sense cDNA was biotin-endlabeled with TdT and probes were hybridized to the Gene 2.0 arrays at 45 °C for 16h with 60 rpms. Hybridised arrays were washed and stained on a Fluidics Station 450 (program: FS450 0002) and scanned on a GeneChip® Scanner 3000 7G (both Affymetrix). Raw image data were analyzed with Affymetrix® Expression Console^TM^ Software (Affymetrix, USA), and gene expression intensities were normalized and summarized with a robust multiarray average algorithm(Irizarry, Hobbs et al. 2003). Transcripts that were expressed differently more than 1.5-fold with a raw p-value lower than 0.05 between the sample groups were categorized as regulated. Enrichment analysis for Wiki pathways was performed using WebGestalt (Wang, Duncan et al. 2013). For the enrichment analysis only, genes changed at least 1.5-fold with a p-value lower than 0.05 between *Foxp3^cre^CREB^fl/fl^* and *CREB^fl/fl^* samples were taken into consideration.

### 2.10 TSDR Methylation analysis

For all methylation analyses, cells from male mice were used. Genomic DNA was prepared from FACS-sorted T_conv_ cells (CD4^+^CD25^-^) as well as T_reg_ cells (CD4^+^CD25^+^Nrp1^+^) from *CREB^fl/fl^* and *Foxp3^cre^CREB^fl/fl^* mice and subsequently converted by bisulfite according to the manufacturer’s instructions (DNeasy Blood & Tissue Kit, Qiagen; EZ DNA Methylation-Lightning Kit, Zymo Research). Pyrosequencing of the Treg cell-specific demethylated region (TSDR) (chromosome position X:7583950-7584149, genome assembly GRCm38.p6) was performed as described previously (Garg, Muschaweckh et al. 2019).

### 2.11 ATAC Seq

Omni-ATAC-seq was performed according to (Corces, Trevino et al. 2017, Li, Schulz et al. 2019) with minor modifications. Prior to transposition, dead cells were removed by centrifugation (800 rpm, 4 min, 4°C). The transposition reaction was with 7.5 μL Tagment DNA Enzyme 1 (TDE1) for 60 ‘min at 37°C. Pre-amplification was with NEBNext Ultra II Q5 Master Mix and Nextera PCR Primers (5 cycles). Quantitative PCR amplification was with NEBNext Ultra II Q5 Master Mix, Nextera PCR Primer and SYBR Gold to determine the number of additional cycles. PCR amplification of additional cycles was as for pre-amplification. PCR fragments were purified with Qiagen MinElute PCR Purification Kit and library concentration and quality were determined by Agilent High Sensitive DNA Kit and TapeStation, respectively. ATACseq libraries were sequenced on the Illumina NextSeq 500 Platform with 75 bps paired end reads in duplicates. ATAC-seq libraries were trimmed with Trim Galore (parameters-q 30 –paired –trim1) and aligned to mouse genome (mm9) using Bowtie2 (Langmead and Salzberg 2012) (parameters-X2000 –no-mixed –no-discordant). Duplicates fragments were removed, and reads were filtered for alignment quality of >Q30 using samtools (Li, Handsaker et al. 2009). Next, we use MAC2 (Zhang, Liu et al. 2008) to perform peak calling (parameters –nomodel –nolambda –keep-dup auto –call-summits). Transcription factor footprinting and differential activity was performed using HINT-ATAC as described in (Li, Schulz et al. 2019).

### 2.12 ELISA

Total IgG and IgE was measured from sera using the the Ready-set-go ELISA system (affymetrix: eBioscience, USA). IL-10 was determined in cell supernatants using Mouse IL-10 ELISA Ready-SET-GO! (2nd Generation), Thermo Fisher Scientific according to manufacturer’s instructions.

### 2.13 Statistical analysis

All data are presented as mean ± standard error (SEM). Data were tested for normality using Shapiro-Wilk Normality test. Differences between two groups were evaluated using two-tailed unpaired or paired (if indicated), Student’s t-test. A two-tailed Mann Whitney test was used, if data were not normally distributed. ONE-way ANOVA was performed if there are more than two groups. Measurements were taken from distinct samples. All statistical analysis and subsequent graphics generation were performed using GraphPad Prism version 7.0 and 8.0 (GraphPad Software, USA). A *p*-value <0.05 was statistically significant.

### 2.14 Study approval

The animal study was approved by the regional government authorities and animal procedures were performed according to German legislation for animal protection. Permission for the projects was granted by the Regierungspräsident/LANUV Nordrhein-Westfalen (81-02.04. 2017.A393).

### 2.15 Data availability

The ATAC-Seq and microarray raw datasets have been deposited in GEO under the accession codes GSE157693 and GSE157933. All data that support the findings of this study are available within the article and Supplementary Files. All other data, including raw data used in each figure will be provided upon reasonable request to the corresponding author.

## 3 RESULTS

### 3.1 CREB enhances Treg numbers but downregulates Foxp3 expression in Tregs

CREB is a transcriptional activator and has been described to critically stabilize Foxp3 expression (Kim and Leonard 2007). It is therefore widely accepted that CREB promotes FoxP3 expression in T_regs_ (Ruan, Kameswaran et al. 2009, Polansky, Schreiber et al. 2010, Zheng, Josefowicz et al. 2010). To verify this hypothesis in vivo, we generated mice with a Foxp3-specific knockout of CREB (*Foxp3^cre^CREB^fl/fl^*). To our surprise, these mice showed enhanced numbers of CD25^+^/Foxp3^+^ positive T_regs_ in thymus (**Fig. 1A and D**), spleen (**1B and D**) and peripheral lymph node (pLN) **(Fig. 1C and D**). Splenic T_regs_ showed no altered expression of Nrp1, PD-1, CTLA4, and ICOS (**Supp Fig. 1A-B**). There was also no convincing different expression of T_h_ lineage transcription factors GATA3 and RORyT and T-bet in CD4^+^Foxp3^+^ cells within different organs (**Supp. Fig. 2A-E)**. However, *Foxp3^cre^CREB^fl/fl^*mice displayed lower Foxp3 expression in T_regs_ on the single cell level (**Fig. 1E and F**) which can be explained by the fact that CREB directly activates the TSDR (Kim and Leonard 2007) (**Fig. 1G**). To verify Cre expression in the Foxp3-Cre mice, we crossed *Foxp3^cre^CREB^fl/fl^*mice with a Cre-inducible reporter mice (*ROSA^R^*^FP^) and examined reporter gene expression in lymphocytes. RFP expression was confined to CD4^+^CD25^+^ cells, which is of importance to exclude cell extrinsic side-effects (**Fig. 1H and I**).

**Figure 1:**
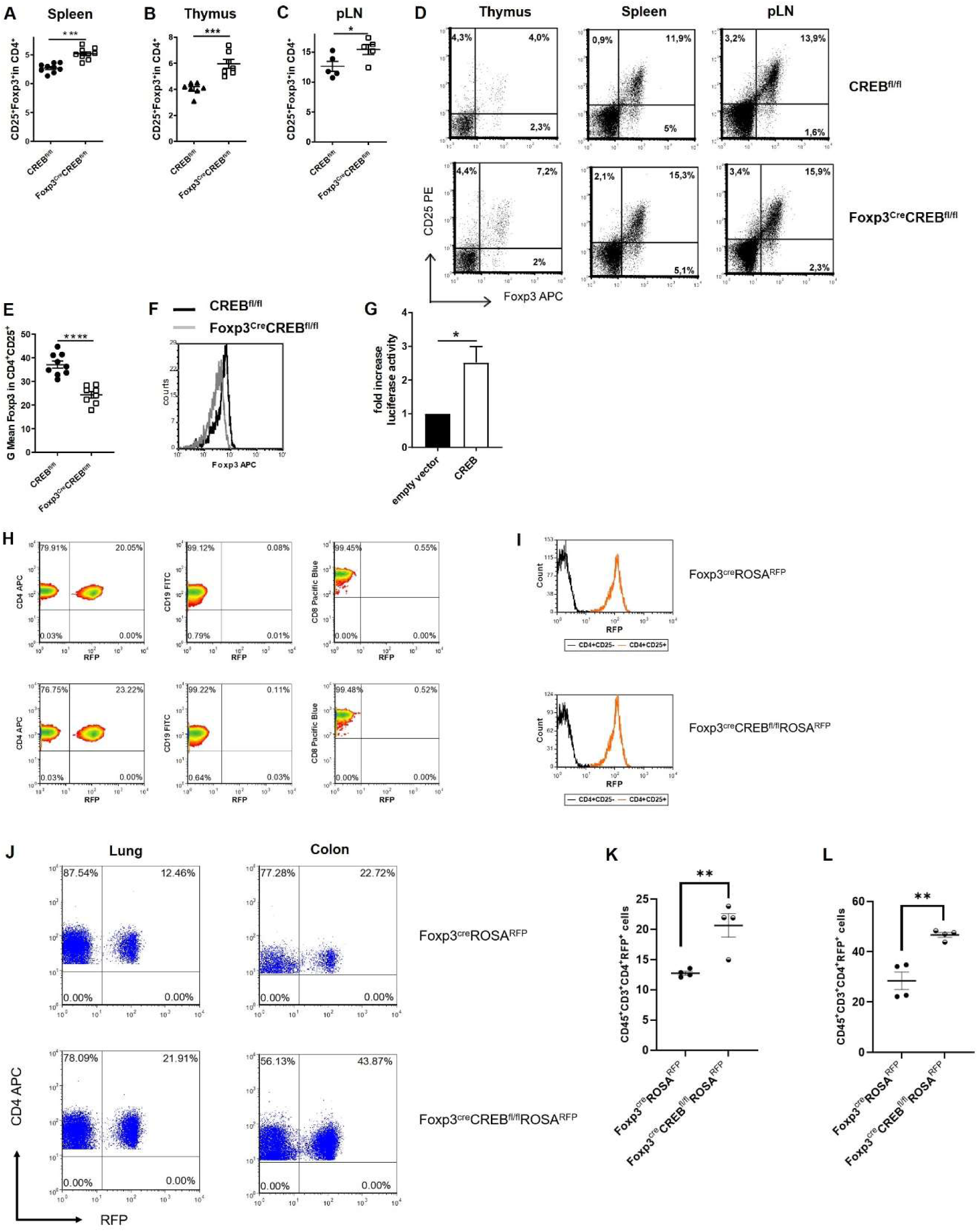
Genetic deletion of CREB in Foxp3^+^ cells enhance percentages of T_regs_ but downregulates their Foxp3 expression. **A)** Statistical analysis of splenic T_reg_ (CD4^+^CD25^+^Foxp3^+^) percentages from *CREB^fl/fl^* and *Foxp3^cre^CREB^fl/fl^* mice (*N* = 9, 4 independently performed experiments, each with 2-3 age and sex-matched mice, 6-9 weeks old). **B)** Statistical analysis of T_reg_ percentages within thymus (CD3^+^CD4^sp^CD25^+^Foxp3^+^) from *CREB^fl/fl^* and *Foxp3^cre^CREB^fl/f^ ^l^*mice (*N* = 7, 3 independently performed experiments, each with 2-3 age and sex-matched mice, 6-9 weeks old). **C)** Statistical analysis of T_reg_ (CD4^+^CD25^+^Foxp3^+^) percentages from pLNs from *CREB^fl/fl^* and *Foxp3^CRE^CREB^fl/fl^*(*N* =5, 2 independently performed experiments, each with 2-3 age and sex-matched mice 7-9 weeks old). **D)** Density plot showing CD25^+^Foxp3^+^ cells gated on CD3^+^CD4^+^CD8^-^ cells in thymus of *CREB^fl/fl^* and *Foxp3^cre^CREB^fl/fl^*mice (left), CD25^+^Foxp3^+^ cells gated on CD4^+^ cells in spleens of *CREB^fl/fl^* and *Foxp3^cre^CREB^fl/fl^* mice (middle) and CD4^+^CD25^+^Foxp3+ cells in pLN of *CREB^fl/fl^* and *Foxp3^cre^CREB^fl/fl^* mice (right). **E**) Statistical analysis of mean fluorescent intensity (MFI) of Foxp3 in *CREB^fl/fl^*and *Foxp3^cre^CREB^fl/fl^* CD4^+^CD25^+^ T cells (*N=6*, 3 independently performed experiments, each with 2 animals, 6-8 weeks old, sex and age matched). **F**) Representative histogram showing overlay of Foxp3 expression in CD4^+^CD25^+^ cells from *CREB^fl/fl^*(black) and *Foxp3^cre^CREB^fl/fl^* (grey) spleens. **G)** RLM cells carrying a stable pGL3-TSDR-FoxPro luciferase Plasmid were transfected either with an empty vector or CREB plasmid, stimulated with PMA and luciferase activity was measured after 4 hours (*N = 5*). **H)** Percentages of RFP^+^ cells within the CD4^+^ (on the left), CD19^+^ (in the middle) and CD8^+^ population (on the right) in spleens of *Foxp3^cre^ROSA^RFP^* and *Foxp3^cre^CREB^fl/fl^ROSA^RFP^*mice. **I)** Representative histogram showing MFI of RFP expression in CD4^+^CD25-(black) and CD4^+^CD25^+^ cells (red) in spleens of *Foxp3^cre^ROSA^RFP^ and Foxp3^cre^CREB^fl/fl^ROSA^RFP^* mice. **J)** Percentages of RFP^+^ cells within the CD45^+^CD3^+^CD4^+^ population in lung and colon of *Foxp3^cre^ROSA^RFP^* and *Foxp3^cre^CREB^fl/fl^ROSA^RFP^* mice. **K)** Statistical analysis of CD45^+^CD3^+^CD4^+^RFP^+^ cells in lung tissue of *Foxp3^CRE^ROSA^RFP^ and Foxp3^cre^CREB^fl/fl^ROSA^RFP^* mice (*N=4*, three independently performed experiments) **L)** Statistical analysis of CD45^+^CD3^+^CD4^+^RFP^+^ cells in colon tissue of *Foxp3^cre^ROSA^RFP^*and *Foxp3^cre^CREB^fl/fl^ROSA^RFP^* mice (*N=4*, three independently performed experiments). A two-tailed, unpaired t-tests was used to test significance. *p<0.05, **p<0.01, ***p<0.001, ****p<0.0001 and results are expressed as the mean ± SEM.

We further used *Foxp3^cre^CREB^fl/fl^ ROSA^R^*^FP^ mice to assess percentages of T_regs_ in non-lymphoid organs. While we did not observe differences in liver tissue, we found a particularly marked upregulation of T_reg_ percentages within lungs and colon of *Foxp3^cre^CREB^fl/fl^ ROSA^R^*^FP^ mice compared to *Foxp3^cre^ROSA*^RFP^ mice (**Fig. 1J-L**) and enhanced absolute numbers of T_regs_ in colon of Foxp*3^cre^CREB^fl/fl^ ROSA^R^*^FP^ (**Supp. Fig. 7B**). This prompted us to analyse if CREB expression in T_regs_ differs in different tissues. We could not find an association of high tissue dependent CREB expression and effects of CREB deficiency on T_reg_ percentages, however found a quite high expression of CREB in murine colon T_regs_ (**Supp. Fig. 8**).

### 3.2 CREB-deficient Tregs are suppressive *in vitro*

To test if lower Foxp3 expression affects suppressive capacity of CREB deficient T_regs,_ we co-cultured *CREB^fl/fl^* T_regs_ and *Foxp3^cre^CREB^fl/fl^* T_regs_ with anti-CD3 and anti-CD28 stimulated WT CD4^+^ cells. Surprisingly, *Foxp3^cre^CREB^fl/fl^* T_regs_ revealed an even enhanced capacity to reduce CD4^+^ T cell proliferation (**Fig. 2A and B**). These data suggest that deletion of CREB, despite reducing Foxp3 expression per cell, enhances function of T_regs_. In addition, expression of Helios was enhanced (**Fig. 2C**) and bisulfite sequencing revealed a demethylated TSDR (**Fig. 2D**), which suggests a stable and suppressive phenotype (Thornton, Lu et al. 2019). In conclusion, despite reduction of Foxp3 levels per cell, CREB deficiency leads to stable and functional T_regs_ *in vitro*.

**Figure 2:**
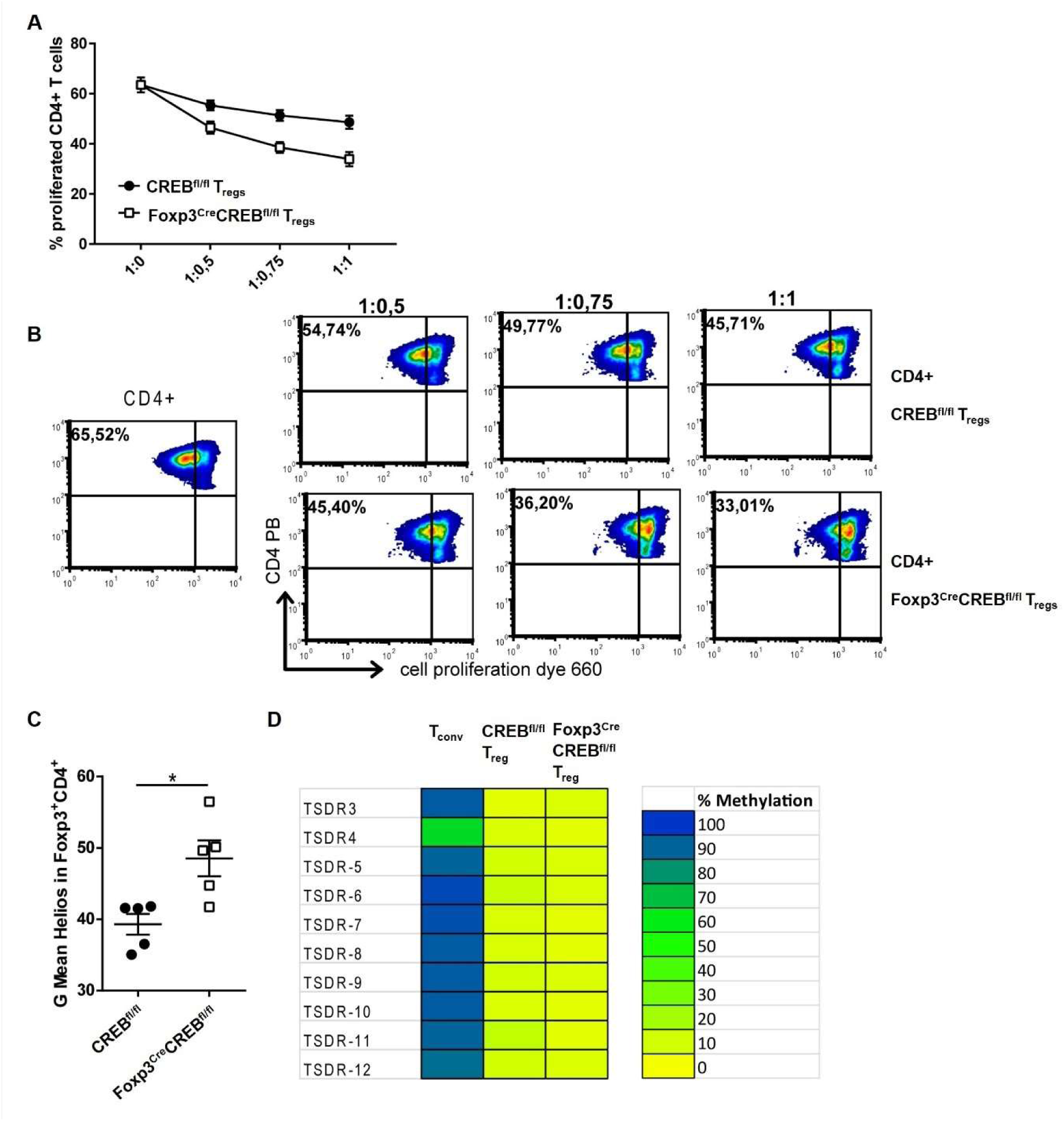
***In vitro* suppressive capacity of T_regs_ is enhanced in the absence of CREB. A)** WT CD4^+^ T cells from spleens (T_eff_) were labeled with efluor660 and stimulated with anti-CD3/CD28 antibodies. CD4^+^CD25^+^ T cells (T_regs_) from *CREB^fl/fl^* or *Foxp3^cre^CREB^fl/fl^* animals were added in different ratios. The proliferation of responder T cells was assessed by efluor660 (ebioscience) dilution (*N = 3*) **B)** Representative density plots of A). **C)** Statistical analysis of MFI of Helios in *CREB^fl/fl^* and Foxp*3^cre^CREB^fl/fl^* CD4^+^Foxp3^+^ T cells (*N = 5*, 2 independently performed experiments, each with 2-3 animals, 8 weeks old sex-matched, a two-tailed Mann Whitney test was performed to test for significance). **D)** Representative TSDR methylation patterns of purified CD4^+^CD25^+^Nrp1^+^ T_reg_ cells and CD4^+^CD25^−^ conventional T cells isolated from spleens of 6-8 weeks old male mice (*N = 4*). Amplicons are vertically arranged, each representing a single CpG-motif. The methylation rate of each motif was translated into a yellow-green-blue color code. *p<0.05 and all results are expressed as the mean ± SEM.

### 3.3 CREB deficient Tregs reveal gene expression signature with enhanced expression of IL-10, IL-4, IL-13 and ST2

To understand why CREB deficient T_regs_ are suppressive despite reduced Foxp3 expression, we performed a whole transcriptome analysis (Affymetrix HTA2 arrays) of flow sorted (CD4^+^CD25^+^Nrp-1^+^) *Foxp3^cre^CREB^fl/fl^* and *CREB^fl/fl^* T_regs_ (**Supp. Fig. 3A**) and appropriate wild-type mice because at the time of this analysis the *Foxp3^CRE^CREB^fl/fl^ ROSA^R^*^FP^ mice were not yet available. Using a fold–change of 1.5 and a p-value of less than 0.05 we found 111 downregulated and 122 upregulated genes in the *Foxp3^cre^CREB^fl/fl^* compared to *CREB^fl/fl^* T_regs_, among those were IL-10 and IL-13 the 2 most strongly upregulated in the *Foxp3^cre^CREB^fl/fl^* T_regs_ (**Fig. 3A)**. KEGG pathway analysis revealed an enrichment of differentially regulated genes in pathways that are associated with Th2 cytokines, such as asthma, Th2 and Th2 cell differentiation and cytokine-cytokine receptor interaction pathways (**Fig. 3B**). Higher *Il13 and Il10* mRNA expression in *Foxp3^cre^CREB^fl/fl^* T_regs_ could be further confirmed by RT-qPCR (**Fig. 3C** and **D**). CD4^+^CD25^+^Foxp3^+^ cells also showed enhanced IL-10 protein levels after 3 days of CD3/CD28 stimulation as assessed by flow cytometry (**Fig. 3E and F**). *In vitro* T_reg_ differentiation with TGF-β revealed a reduced capacity of *Foxp3^cre^CREB^fl/fl^* T cells to differentiate towards Foxp3^+^ cells, while IL-10 expressing cells within Foxp3^+^ cells were enhanced. Furthermore IL-10 cytokine expression of TGF-β stimulated *Foxp3^cre^CREB^fl/fl^* T cells measured by ELISA was enhanced as well as *Il13, Il5 and Il10* mRNA expression, which suggests that CREB deficiency in T_regs_ induces a Th2 biased phenotype (**Fig. 3H-L**). Beyond cytokines, expression of Il1r1 (ST2) showed enhanced expression on the microarray. ST2, the receptor of IL-33 is preferentially expressed on colonic T_regs_, where it promotes T_reg_ function and adaption to the inflammatory environment. It provides a necessary signal for T_reg_-cell accumulation and maintenance in inflamed tissues, and it was shown to regulate IL-10 expressing T_regs_ in the gut (Schiering, Krausgruber et al. 2014). In addition, IL-33 has been identified as a major interleukin in asthma pathology (Drake and Kita 2017). ST2 was highly expressed in the *Foxp3^cre^CREB^fl/fl^* T_regs_, which was confirmed by flow cytometry (**Fig. 3M and N**). Moreover, the whole transcriptome analysis also revealed an upregulated expression of CREM in *Foxp3^cre^CREB^fl/fl^* T_regs_ and qRT-PCR also showed tendentially enhanced CREM expression in *Foxp3^cre^CREB^fl/fl^* T_regs_ **(Fig. 3O**). In conclusion, these data suggest that CREB deletion induces ST2^+^ T_regs_. This finding led us to investigate the protein expression of ST2 within different organs in *Foxp3^cre^ROSA^RFP^*and *Foxp3^cre^CREB^fl/fl^ROSA^RFP^*mice. ST2 expression was clearly enhanced in T_regs_ in the thymus, spleen, mesenteric lymph node, lung and colon of these mice **(Fig. 3P-T)**.

**Figure 3:**
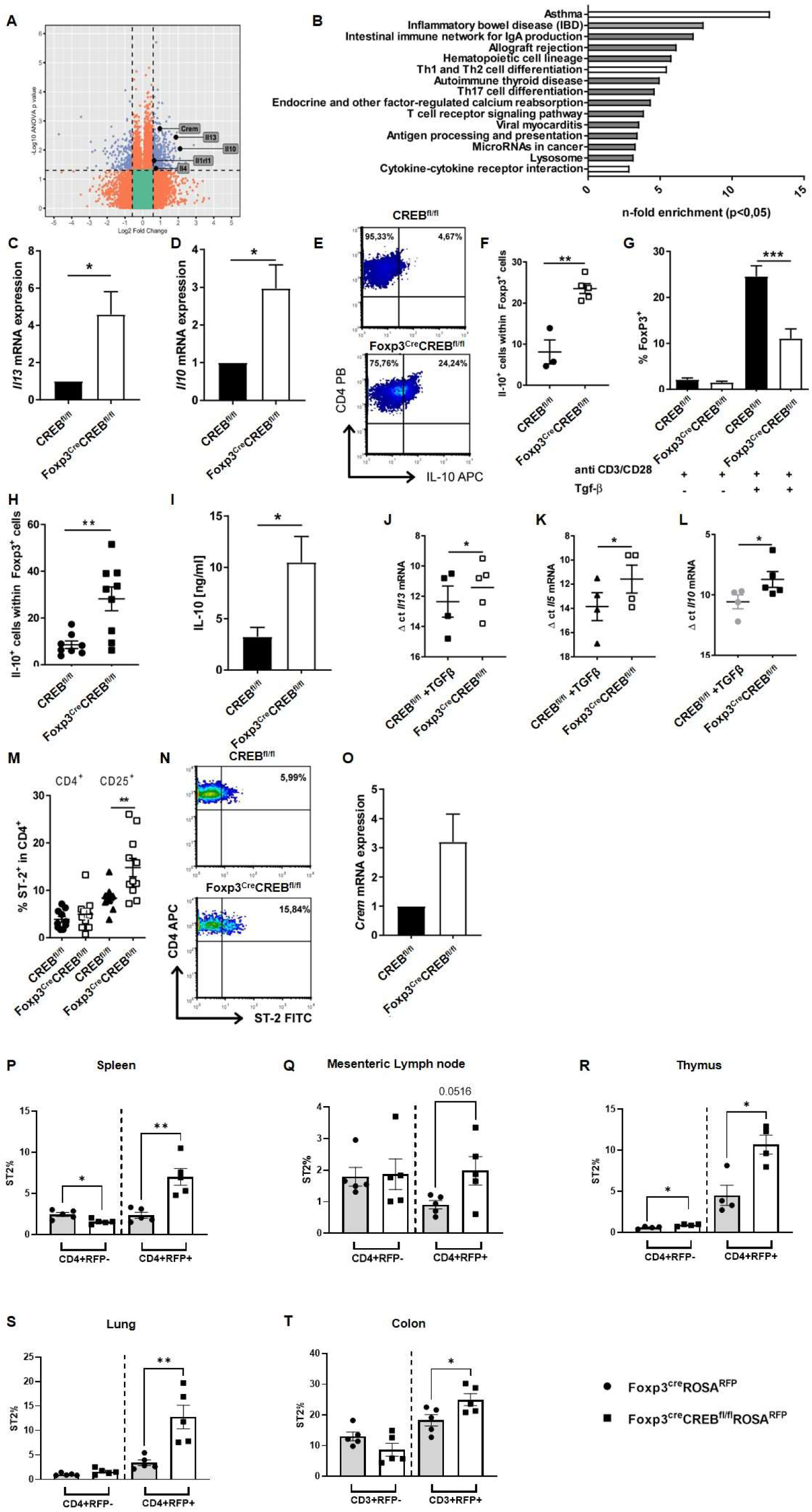
Altered gene expression in CREB deficient T_regs_. **A)** Gene expression in T_regs_ from spleens of *CREB^fl/fl^* and *Foxp3^cre^CREB^fl/fl^ mice.* Colors indicate significant upregulation or downregulation of at least 1.5-fold with p-value not more than 0.05 (blue); regulation of at least 1.5-fold or regulation with p-value not more than 0.05 (red). **B)** Selection of pathways and associated genes, which were significantly enriched. **C)** N-fold *Il13* mRNA expression in splenic T_regs_ from from *CREB^fl/fl^* and *Foxp3^cre^CREB^fl/fl^* mice (*N = 5*, male, 8-10 weeks old) analyzed by RT-qPCR. Bars indicate mean and error bars SEM, two-tailed one-sample test. **D)** N-fold *Il10* mRNA expression in splenic T_regs_ from *CREB^fl/fl^* and *Foxp3^cre^CREB^fl/fl^* mice (*N = 5*, male, 8-10 weeks old) analyzed by RT-qPCR. Bars indicate mean and error bars SEM, two-tailed one sample test. **E)** MACS-isolated T cells from spleens were stimulated ex-vivo with anti-CD3/CD28 antibodies for three days and re-stimulated with PMA/Ionomycin in the presence of Golgi Plug^TM^. A representative density plot shows IL-10 production in CD4^+^Foxp3^+^ cells **F)** Statistical analysis of IL-10 expression of Foxp3^+^ cells after treatment as in E) (*N = 3* independent experiments, each dot represents one animal, two tailed unpaired t-test. **G)** CD4^+^CD25^-^ T cells were stimulated with anti-CD3/CD28 antibodies in the presence/absence of 5 ng/ml TGF-β for five days. Percentages of Foxp3^+^ cells from 6 independently performed experiments were determined, each with spleen from one mouse (*N =13 CREB^fl/fl^* and *N=12 Foxp3^cre^CREB^fl/fl^* mice), a two tailed unpaired t-test was used to test for significance. **H)** Statistical analysis of IL-10^+^ cells within Foxp3^+^ cells after treatment as in G). An unpaired, two tailed t-test was used to test for significance, each dot represents one animal and error bars SEM **I)** IL-10 in supernatants of CD4^+^CD25^-^ T cells after stimulation with anti-CD3/CD28 antibodies in the presence of 5 ng/ml TGF-β for five days measured by ELISA, an unpaired two-tailed t test was performed to test for significance, bars indicate mean and error bars SEM (*N = 6*). **J)-L)** Δ ct mRNA expression of cells after treatment as in G, each dot represents one animal and error bars SEM, two tailed unpaired t-tests were used to test for significance. **M)** Statistical analysis of splenic ST2^+^ cells within CD4^+^ and CD4^+^CD25^+^ cells of *CREB^fl/fl^*and *Foxp3^cre^CREB^fl/fl^* mice. 3 independently performed experiments (*N = 10 CREB^fl/fl^* and *N =11 Foxp3^cre^CREB^fl/fl^* mice), a two tailed unpaired t-test was used to test for significance, each dot represents one animal and error bars SEM. **N)** Representative density plot of M. **O)** N-fold *Crem* mRNA expression in splenic T_regs_ from from *CREB^fl/fl^* and *Foxp3^cre^CREB^fl/fl^*mice (N = 3, male, 8-10 weeks old) cells analyzed by RT-qPCR. **P-T)** Statistical analysis of ST2^+^ in CD4+ and CD4+RFP+ cells in *Foxp3^cre^ROSA^RFP^* (*N = 5*) and *Foxp3^cre^CREB^fl/fl^ROSA^RFP^* (N = 5) in **P)** spleen, **Q)** mesenteric lymph node, **R)** thymus, **S)** lung, bars indicate mean and error bars SEM, two tailed unpaired t-test was used to test for significance. **T)** Statistical analysis of ST2+ in CD3+ and CD3+RFP+ cells in *Foxp3^cre^ROSA^RFP^* (*N = 5*) and *Foxp3^cre^CREB^fl/fl^ROSA^RFP^* (*N = 5*) in colon, two tailed unpaired t-test was used to test for significance. *p<0.05, **p<0.01, ***p<0.001, and results are expressed as the mean ± SEM.

### 3.4 *Foxp3^cre^CREB^fl/fl^*T cells dampen inflammation in experimental colitis

Since ST2^+^T_regs_ also play a protective role in intestinal mucosal inflammation (Schiering, Krausgruber et al. 2014), we performed a transfer of *Foxp3^cre^CREB^fl/fl^* T cells (CD4+CD25-) or appropriate *CREB^fl/fl^* T cells (CD4+CD25-) into lymphopenic Rag2^-/-^ mice. This usually results into development of colitis within 5 weeks. Strikingly, transfer of *Foxp3^cre^CREB^fl/fl^* T cells did not result in the expected weight loss (**Fig. 4A**) and recipient mice displayed decreased intestinal inflammation (**Fig. 4B**-**C**). Furthermore, inflammatory cytokines in the gut were markedly reduced (**Fig. 4D**) as well as IL-17 and IFN-γ production in CD4^+^ T cells in the mesenteric lymph nodes (mLNs) (**Fig. 4E**-**G**). The experiment was repeated with CD4+CD25-T cells of *Foxp3^cre^ROSA^R^*^FP^ and *Foxp3^cre^CREB^fl/fl^ ROSA^R^*^FP^ mice into Rag2-/-mice and ending with the same results. The transfer of CD4+CD25-T cells of *Foxp3^cre^CREB^fl/fl^ ROSA^R^*^FP^ mice prevented development of colitis and resulted in enhanced accumulation of RFP+ T_regs_ in different organs (**Fig. 5A-G**). Interestingly, RFP+ T_regs_ also accumulated in the liver, while we did not find a difference in the RFP expression in the *Foxp3^cre^CREB^fl/fl^ ROSA^R^*^FP^ mice under steady state conditions.

**Figure 4:**
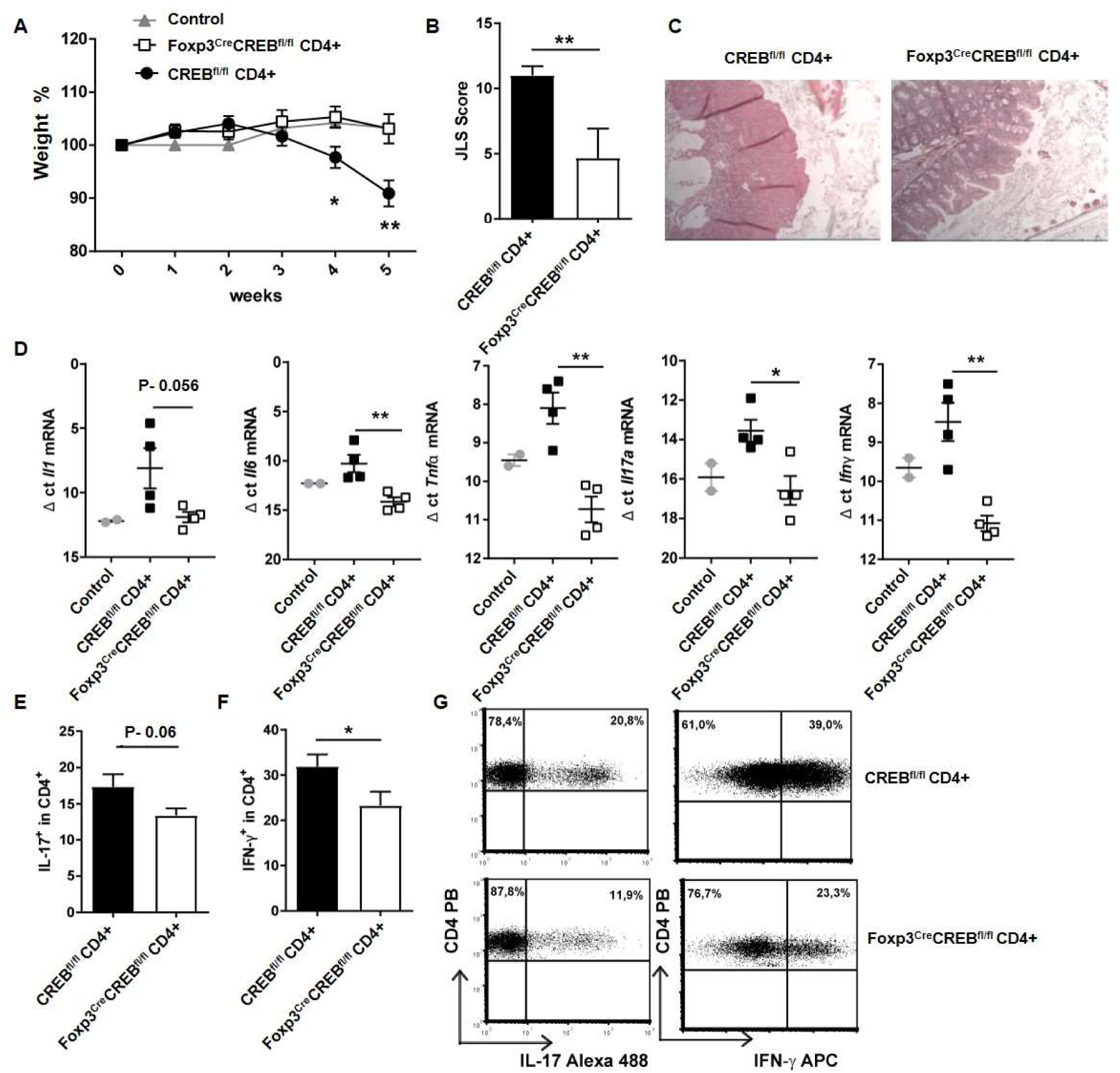
***Foxp3^cre^CREB^fl/fl^*CD4+ T cells induce only moderate inflammation in experimental colitis.** Rag2^-/-^ mice were adoptively transferred with CD4^+^ CD25^-^ wild-type cells (*CREB^fl/fl^* CD4^+^) or *Foxp3^cre^CREB^fl/fl^* CD4^+^ CD25^-^ T cells. Untreated Rag^-/-^ mice were used as controls. Mice were weighed and sacrificed 5 weeks after transfer. **A)** Body weight as a percent of starting weight, two-tailed unpaired t-test were used to test for significance (*N = 11 CREB^fl/fl^*, *N = 9 Foxp3^cre^CREB^fl/fl^* recipients and *5* control Rag^-/-^ mice were analyzed in 3 independently performed experiments). **B)** Results of histological JLS (The Jackson Laboratory Scoring) score of colon sections (two independently performed experiments, *CREB^fl/fl^* recipients N = 5, *Foxp3^cre^CREB_fl/fl_* recipients N = 6, a two-tailed Mann Whitney test was used). **C)** Representative photomicrographs of hematoxylin and eosin (H&E)-stained colon sections imaged using a 10x objective. **D)** Expression of inflammatory cytokines analyzed by RT-qPCR. Dots represent Δ ct values normalized to β-actin. Each dot represents one animal. (ct levels are inversely proportional to the amount of target nucleic acid in the sample), two tailed unpaired t-tests were used to test for significance, each dot represents one animal (*N = 2* control mice, *N = 4 CREB^fl/f^*, *N = 4 Foxp3^cre^CREB^fl/fl^* recipients). **E)** Statistical analysis of IL-17^+^ cells within CD4^+^ cells in mLNs (three independently performed experiments, *CREB^fl/fl^* recipients N = 9, *Foxp3^cre^CREB^fl/fl^*recipients N = 9), a two-tailed, unpaired t-test was used. **F)** Statistical analysis of IFN-γ^+^ cells within CD4^+^ cells in mLNs. (three independently performed experiments, *CREB^fl/fl^* recipients *N = 11*, *Foxp3^cre^CREB^fl/fl^*recipients *N = 9*), two-tailed, unpaired t-test was used. **G)** Representative dot plots of E and F. *p<0.05, **p<0.01 and results are expressed as the mean ± SEM.

**Figure 5:**
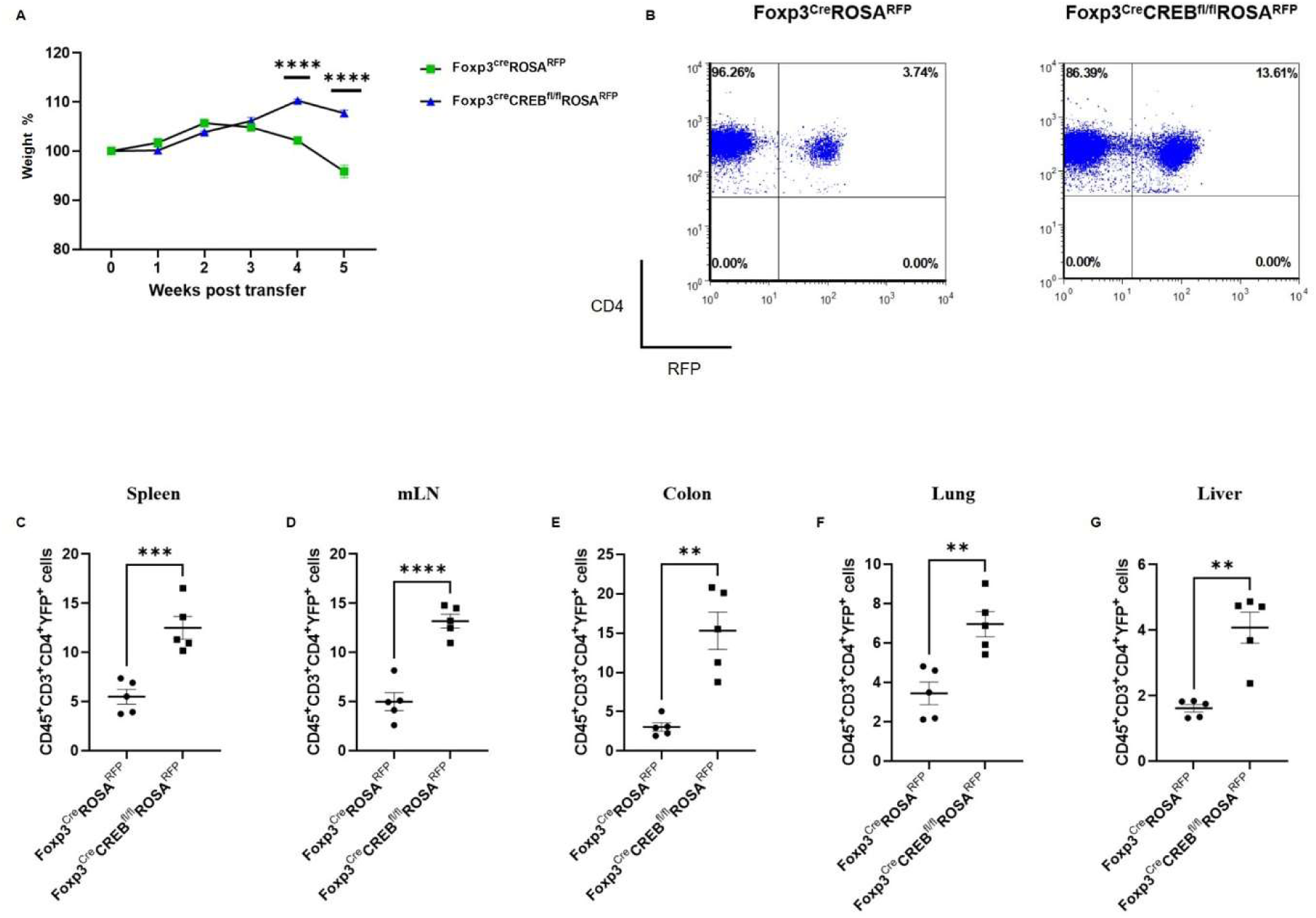
*Foxp3^cre^CREB^fl/fl^ROSA^RFP^*CD4+ T cells induce only moderate inflammation in experimental colitis, while CD4+RFP+ cells expand in peripheral tissues. Rag2^-/-^ mice were adoptively transferred with CD4^+^ CD25^-^ wild-type cells (*Foxp^cre^ROSA^RFP^*CD4^+^) or *Foxp3^cre^CREB^fl/lf^ROSA^RFP^* CD4^+^ CD25^-^ T cells. Mice were weighed and sacrificed 5 weeks after transfer. **A)** Body weight as a percent of starting weight, two-tailed unpaired t-test were used to test for significance (*N = 5 Foxp^cre^ROSA^RFP^* and *N = 5 Foxp3^cre^CREB^fl/lf^ROSA^RFP^* recipient mice were analyzed in 1 independently performed experiment). **B)** Representative Flow cytometry gated on CD4 and RFP in splenocytes. Percentages of CD45^+^CD3^+^CD4^+^RFP^+^ cells in **C)** Spleen, **D)** mLNs, **E)** Colon, **F)** Lung, and **G)** Liver, two-tailed, unpaired t-test was used. **p<0.01, ***p<0.001, ****p<0.0001 and results are expressed as the mean ± SEM.

### 3.5 Blockade of IL-10 but not of ST2 signalling reverses protective effects of *Foxp3^cre^CREB^fl/fl^* T cells in experimental colitis

ST2^+^ T_regs_ are protective in intestinal inflammation (Schiering, Krausgruber et al. 2014), in addition they are Th_2_ biased and release IL-10 (Siede, Frohlich et al. 2016). To decipher if *Foxp3^cre^CREB^fl/fl^*T cells prevent colitis by IL-10 production and/or ST2 expression, we performed an adoptive transfer colitis with anti-IL10R antibodies or recombinant soluble ST2 (sST2). sST2 acts as a decoy for IL-33 and inhibits its biological activity (Holgado, Braun et al. 2019). Blockade of IL-10 signalling induced a colitis in the formerly resistant *Foxp3^cre^CREB^fl/fl^*T cell recipient mice (**Fig. 6A-C**), while ST2 blockade had no effect on disease progression (**Supp. Fig. 4**).

**Figure 6:**
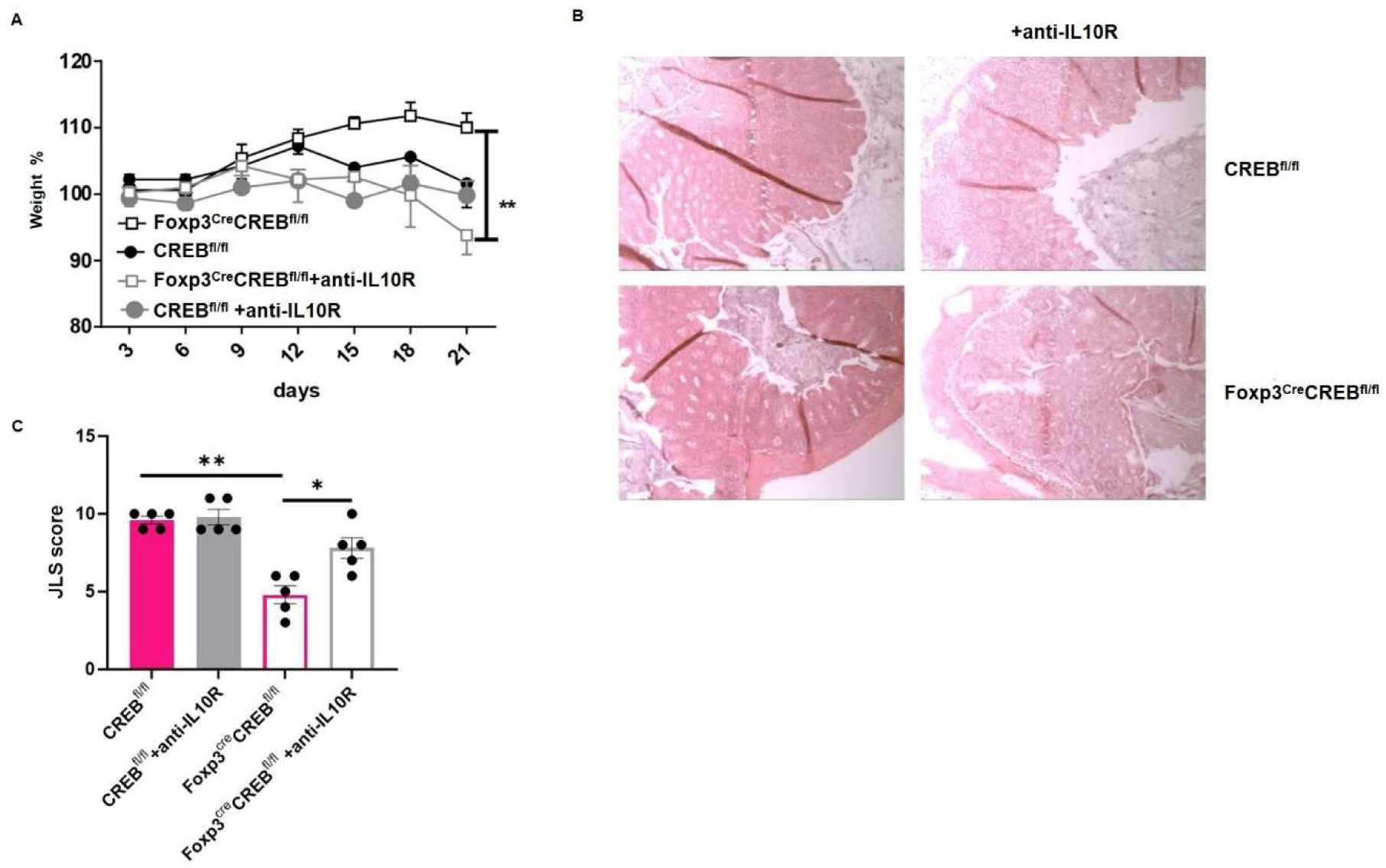
**Reduced colitis in *Foxp3^cre^CREB^fl/fl^* CD4^+^ T cell recipients depends on IL-10 signaling**. Rag2^-/-^ mice were adoptively transferred with CD4^+^ wild-type cells (WT CD4^+^) or *Foxp3^cre^CREB^fl/fl^* CD4^+^ T cells. Mice were either treated with an anti-IL10R antibody or an isotype control. **A)** Body weight as a percent of starting weight, Bars indicate mean and error bars SEM (*N=5* animals in each group, a two-tailed unpaired t-test was used, two independently performed experiments). **B)** Representative photomicrographs of a hematoxylin and eosin (H&E)-stained colon section from *CREB^fl/fl^*and *Foxp3^cre^CREB^fl/fl^* CD4^+^ T cell recipients, that were treated with anti-IL10R, imaged using a 10x objective. **C)** Results of histological JLS (The Jackson Laboratory Scoring) score of colon sections (two independently performed experiments with overall *N=5* mice in each group, two-independently performed experiments, a two tailed Mann Whitney test was used to test for significance). *p<0.05, **p<0.01 and results are expressed as the mean ± SEM.

### 3.6 Genome-wide chromatin accessibility of WT and CREB-deficient Tregs

To analyse if CREB deletion alters chromatin accessibility at different promoters and enhancers, we performed the Assay for Transposase-Accessible Chromatin using sequencing (ATAC-seq) in *CREB^fl/fl^* and *Foxp3^cre^CREB^fl/fl^* T_regs_. We observed a decrease in open chromatin around CREB and CREM motifs on *Foxp3^cre^CREB^fl/fl^* mice, which indicate loss of binding activity of these transcription factors in these cells (**Fig. 7A**). The *Il1rl1* (ST2) region showed significant differences in open chromatin around the promoter and potential enhancer sites in T_regs_ of *Foxp3^cre^CREB^fl/fl^* mice, while CREB-binding motifs could not be identified (**Fig. 7B**). In addition, the IL-13 enhancer region showed clear differences between the 2 mice strains. Within this region, binding sites for Nrf1 were identified, which is a CREB dependent transcription factor (Suliman, Sweeney et al. 2010). Moreover, we found a change in open chromatin of an alternate promoter of CREM (**Fig. 7B**). CREM has been identified as one of the most regulated genes in ST-2 positive colonic T_regs_ of healthy mice before (Miragaia, Gomes et al. 2019). To prove if CREM is involved in the regulation of ST2 we generated *Foxp3^cre^CREB^fl/fl^CREM^-/-^*mice. *In vitro* differentiation towards Foxp3+ cells was not significantly diminished in *Foxp3^cre^CREB^fl/fl^CREM^-/-^* mice (**Fig. 7C**). Furthermore, deletion of CREM normalized Foxp3 expression per cell in CREB-deficient T_regs_ at least to some extent (**Fig. 7D**), however reversed expression of ST2 (**Fig. 7E**). Taken together our data suggest that the interaction of CREB and CREM determine the expression of Foxp3 and ST2 and induction of CREM in CREB deficient T_regs_ contributes to reduced Foxp3 expression. To prove the effect in vivo, we performed a transfer of *Foxp3^cre^CREB^fl/fl^*CREM^-/-^ T cells (CD4+CD25-) or appropriate wild type T cells (CD4+CD25-) into lymphopenic Rag2^-/-^ mice. The additional genetic deletion of CREM reversed the effect of the CREB deletion and resulted in enhanced colitis severity measured by JLS (The Jackson Laboratory Scoring) score in the transferred mice (**Fig. 7F**).

**Figure 7:**
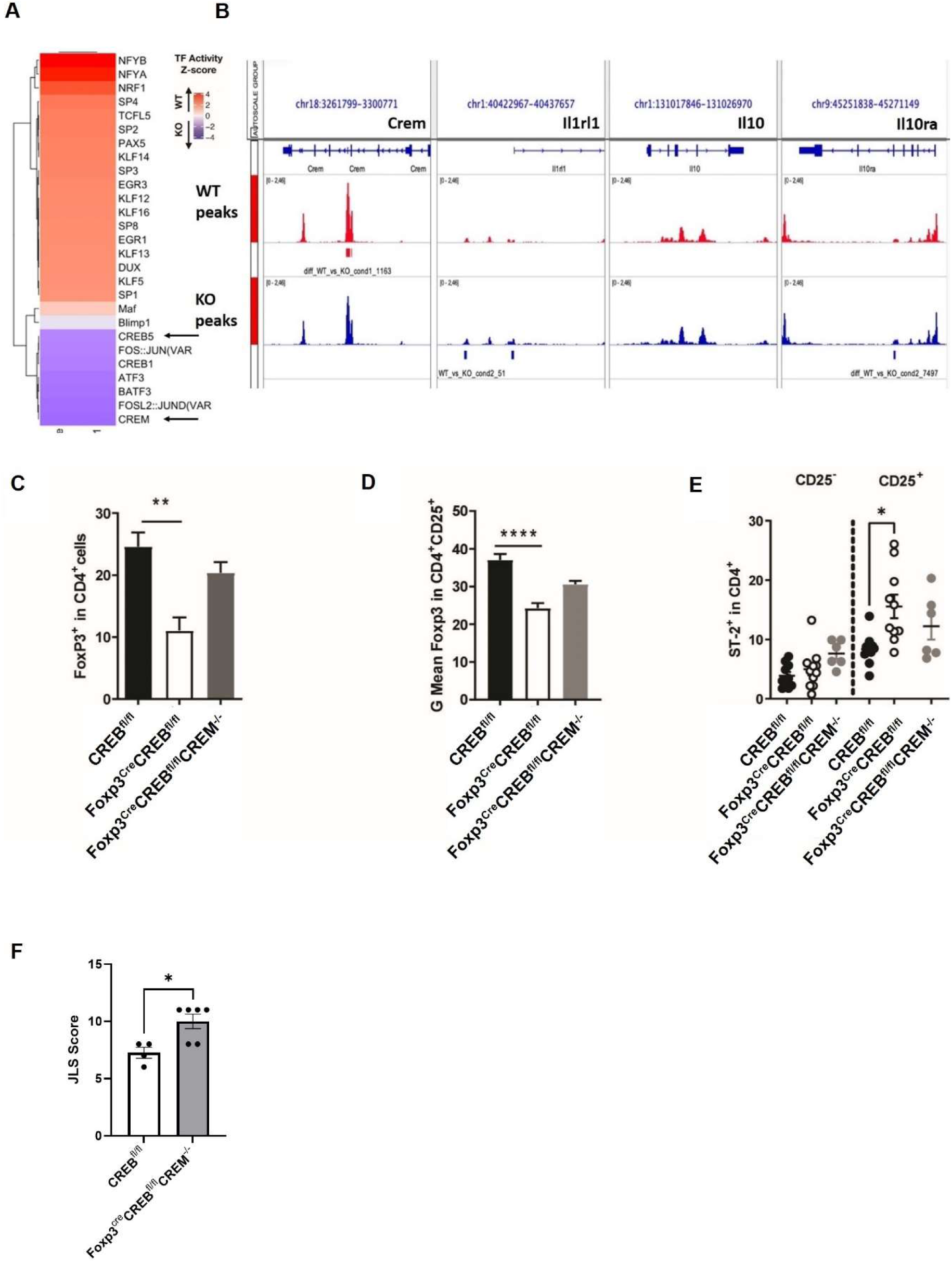
Genome-wide chromatin accessibility of *CREB^fl/fl^* and *Foxp3^cre^CREB^fl/fl^* mice. **A)** ATAC-seq foot printing analysis indicates decrease in TF activity for CREB and CREM upon CREB KO and increase in TF activity of NRF1 upon CREB KO (p-value < 0.05; z-test). **B)** Example of genes with loss of chromatin accessibility upon CREB KO includes an alternative promoter of CREM (red peaks) and enhancer regions around SC-2/Il1nl1 (blue peaks) (adjusted p-value < 0.05; MACS2 diff. peak caller). **C)** CD4^+^CD25^-^ T cells were stimulated with anti-CD3/CD28 antibodies in the presence/absence of 5 ng/ml TGF-β for five days. Percentages of Foxp3^+^ cells from 6 independently performed experiments were determined, each with spleen from one mouse, *N* = 13 *CREB^fl/fl^*, *N = 12 Foxp3^cre^CREB^fl/fl,^* and *N = 3 Foxp3^cre^CREB^fl/fl^CREM*^-/-^ mice, a two tailed unpaired t-test was used to test for significance, bars indicate mean and error bars SEM. **D)** Statistical analysis of MFI of Foxp3 in *CREB^fl/fl^* (*N* = 13) and *Foxp3^cre^CREB^fl/fl^* (N = 9) and *Foxp3^cre^CREB^fl/fl^CREM*^-/-^ (*N* =3) CD4^+^CD25^+^ T cells, bars indicate mean and error bars SEM, two tailed unpaired t-test was used to test for significance. **E)** Statistical analysis of ST2^+^ in CD4^+^ and CD4^+^CD25^+^ cells in *CREB^fl/fl^* (*N* = 5) and *Foxp3^cre^CREB^fl/fl^*(*N = 6*) and *Foxp3^cre^CREB^fl/fl^CREM*^-/-^ (*N* =3) CD4^+^CD25^+^ T cells, bars indicate mean and error bars SEM, one-way Anova test was used to test for significance. **F)** Rag2^-/-^ mice were adoptively transferred with *CREB^fl/fl^* CD4 T-cells (CD4^+^CD25-) or *Foxp3^cre^CREB^fl/fl^* CREM^-/-^ CD4^+^ T cells (CD4+CD25-). Results of histological JLS (The Jackson Laboratory Scoring) score of colon sections (one experiment with overall *N=4/5* mice in each group, a two tailed Mann Whitney test was used to test for significance). *p<0.05, **p<0.01, ****p<0.0001 and results are expressed as the mean ± SEM.

## 4 DISCUSSION

In this study we provide evidence that CREB cell-intrinsically regulates Foxp3 expression in T_regs_ and thereby plays a critical role in T_reg_ cell-mediated immune homeostasis. In detail, CREB deficient T_regs_ show a reduced expression of Foxp3 per cell but enhanced expression of ST2, IL-10, and IL-13. Enhanced IL-10 expression of CREB-deficient T_regs_ prevents T cell mediated colitis. Enhanced IL-10 expression is also a hallmark of ST2^+^ T_regs_. ST2 is expressed on colonic T_regs_ and enhances local intestinal T_reg_ cell responses (Schiering, Krausgruber et al. 2014). We identified particularly enhanced expression of T_regs_ in colon and lung in *Foxp3^cre^CREB^fl/fl^* mice, which also includes upregulation of ST2 expression, and which might contribute to the unexpectedly protective role of CREB deletion in T_regs_ in type I immune responses. This protective role is in line with Wang et al., who demonstrated an important role for CREB in regulating the balance between Th17 cells and T_regs_ (Wang, Ni et al. 2017). They showed that mice bearing a deletion of CREB in all CD4^+^ T cells prevented T cell mediated transfer colitis comparable to our model. Moreover, these mice were also resistant in an EAE model. This protective effect in EAE was surprisingly lost, when CREB was specifically deleted in T_regs_. Nevertheless, except from the EAE model Wang et al. did not analyse the Foxp3 specific CREB deleted mice in further detail.

We hypothesize that in particular the enhanced IL-10 secretion prevented colitis in Rag-/-mice transferred with *Foxp3^cre^CREB^fl/fl^*CD4+ T cells. IL-10 is an important regulator of intestinal homeostasis, as IL10 and IL10R deficient mice spontaneously develop intestinal inflammation (Kuhn, Lohler et al. 1993, Spencer, Di Marco et al. 1998) and blocking IL-10 signalling induced colitis in *Foxp3^cre^CREB^fl/fl^* T cell recipients that were previously resistant. It has been shown before that CREB induces IL-10 transcription in macrophages. Our data thus were highly unexpected and do not support these findings but might instead argue for cell dependent mechanisms that regulate IL-10 expression. Our ATACseq data however do not show a direct effect of CREB expression on chromatin accessibility within the IL-10 enhancer and promoter region, thus we currently cannot mechanistically explain these findings.

We also found enhanced percentages of T_regs_ within the thymus, spleen and pLNs but reduced Foxp3 expression per cell within peripheral lymphoid organs. Wang et al. also suggest that compensation by related transcription factors such as CREM exist. Interestingly and corroborating that theory, we found a clear upregulation of CREM in CREB deficient T_regs_ and additional deletion of CREM recuperates Foxp3 expression in CREB deficient T_reg_ cells, while it downregulates ST2 expression. Strikingly, CREM expression, which was enhanced in *Foxp3^cre^CREB^fl/fl^*T_regs_ was found before as a highly regulated gene in ST2 positive colon and skin T_regs_ (Miragaia, Gomes et al. 2019). Delacher et al. characterized ST2^+^ tissue T_regs_ that are present in all non-lymphoid tissues, which were found to express killer cell lectin-like receptor subfamily G1 (KLRG1) and Th2 associated factors, including IL-10 (Delacher, Imbusch et al. 2017, Delacher, Imbusch et al. 2020). These “tisT_regs_ST-2” cells are programmed by Basic leucine zipper transcription factor ATF-like (BATF) (Delacher, Imbusch et al. 2020) in lymphoid organs and perform important tissue homeostasis and regenerative functions. Our data point to the fact that the expression of CREB and CREM also regulate these tisT_reg_ST2 in non-lymphoid tissues like colon, lung and liver. Spath et al. vigorously analysed ST2 trajectories in mice and showed a high plasticity in ST2 expression and hierarchies in tissue-specific phenotypes (Spath, Roan et al. 2022). In addition, they found that ST2^+^ T_regs_ were highly proliferative and had a high migratory potential. This goes in line with our findings that these cells can be found in spleen, lung, liver, mLN and the colon. They additionally found a high prevalence in skin and VAT, which was not analysed in our setting. When comparing the transcriptional profiles with our analyses we can confirm an upregulation of PTPN13 in our data (1.97fold) and a regulation of Rab4a (-1.51fold), between the CREB deficient T_regs_ and wild type T_regs_, the other factors that distinguished ST2^+^ from ST2^-^ T_regs_ (Lyn, Gata3, Rln3, Klrg1, and Tbc1d4) were not different. Interestingly, in their analysis, ST2^+^ T_regs_ were enriched in more activated, differentiated T_reg_ populations such as ID2^+^ T_regs_ and ID2 was upregulated in our analysis as well (1.7fold) (Spath, Roan et al. 2022).

Limitations of our study are that we mechanistically mainly analysed splenic T_regs_, while ST2 is a hallmark of tissue T_regs_ and further analysis will show how this will affect local T_regs_ and tissue homeostasis. Another issue is that it is not clear how CREB activation is regulated in T_regs_. CREB has nine serine residues in the kinase inducible domain (KID) that can be phosphorylated and activated by different kinases and different phosphorylation patterns of CREB can exert opposite effects. Further research will therefore also be necessary to identify therapeutic targets to directly influence CREB activity in T_regs_. In conclusion, we provide evidence that CREB plays a pivotal role in T_regs_. CREB is an important transcription factor to maintain Foxp3 expression and the interaction of CREM and CREB are crucial for the expression of ST2 in T_regs_. A lack of CREB enhances IL-10 expression and thereby prevents Th1 mediated diseases.

## AUTHOR CONTRIBUTIONS

S.H.S conducted the experiments, analysed the data and wrote the manuscript. J.H, S.S, T.L, T.G, C.N, T.C, M.S, E.V, S.B, K.O & S.F conducted experiments, M.K contributed to study design, K.O, I.C, Z.L, L.G, B.D, & S.F analysed data, J.H & M.Z analysed data and contributed to study design, E.S, R.B, & B.L provided reagents, A.S provided the mice, T.B provided mice and contributed to study design, N.W contributed to reviewing the manuscript, K.O & K.T revised the manuscript and designed the study.

## DECLARATION OF COMPETING INTEREST

The authors declare that they have no known competing financial interests or personal relationships that could have appeared to influence the work reported in this paper.

## ACKNOWLEDGMENTS

This research project was supported by DFG grant TE339/20-1 to K.T. J.H and T.B were supported by the DFG grant SFB/TRR 355/1 (Project number: 490846870). The authors acknowledge the IZKF Flow Cytometry Facility for the use of cell sorter (DFG grant project ID 439895892) and Genomics facility for their service.

± SEM.

APPENDIX A

## Notes

### Competing Interest Statement

The authors have declared no competing interest.

### Summary of Updates

The revised manuscript has been refined to focus specifically on the impact of CREB deletion in regulatory T cells (Tregs) in relation to colitis. This updated version includes new data from adoptive transfer colitis experiments using Foxp3creCREBfl/flROSARFP mice, providing a more comprehensive analysis of CREB's role in Treg function during intestinal inflammation. Supplemental figures have also been added to support the main findings and offer further insights into the mechanisms involved. Additionally, the author affiliations have been updated, reflecting any changes in institutional associations. Overall, the manuscript offers a targeted exploration of the molecular mechanisms underlying Treg-mediated suppression of colitis, potentially providing valuable insights into therapeutic strategies for inflammatory bowel diseases.

