## Supplementary Figures for "CREB REGULATES FOXP3^+^ST-2^+^ TREGS WITH ENHANCED IL-10 PRODUCTION"

### APPENDIX B

#### SUPPLEMENTARY FIGURES

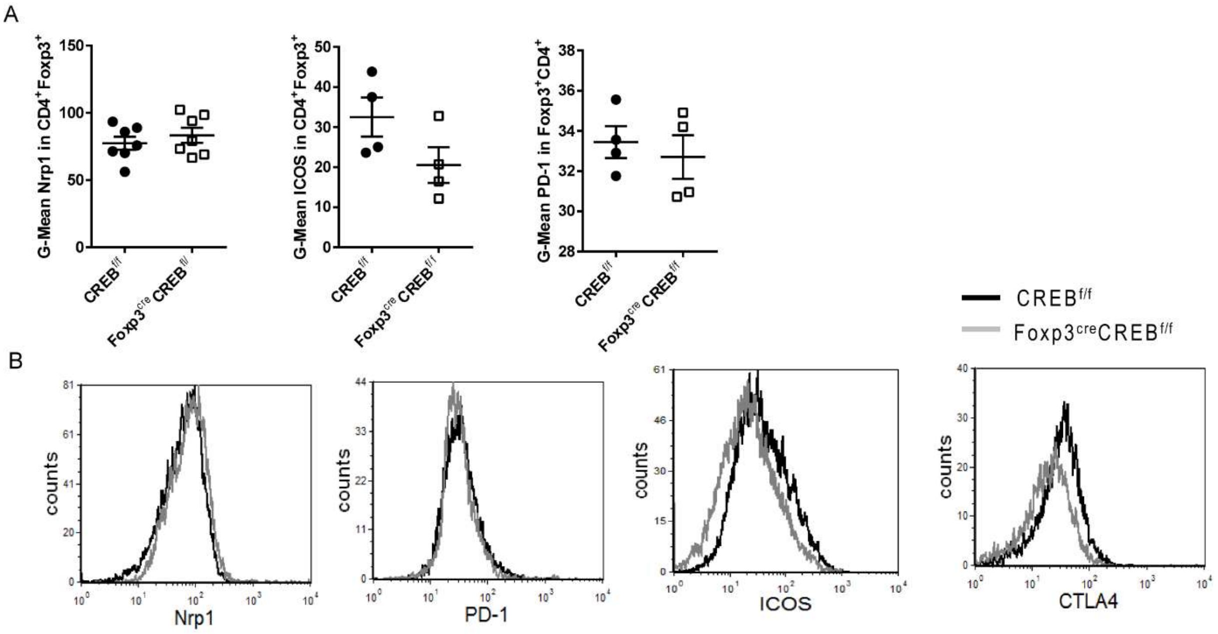


**Supplementary Figure 1: Characterization of CREB-deficient $T_{reg}$ cells. A)** Statistical analysis and **B)** representative histograms of $T_{reg}$ cell markers in $CD4^{+}Foxp3^{+}$ splenic cells. Mice were 6-9 weeks old, and sex and age matched. For A, results are expressed as the mean ± SEM.

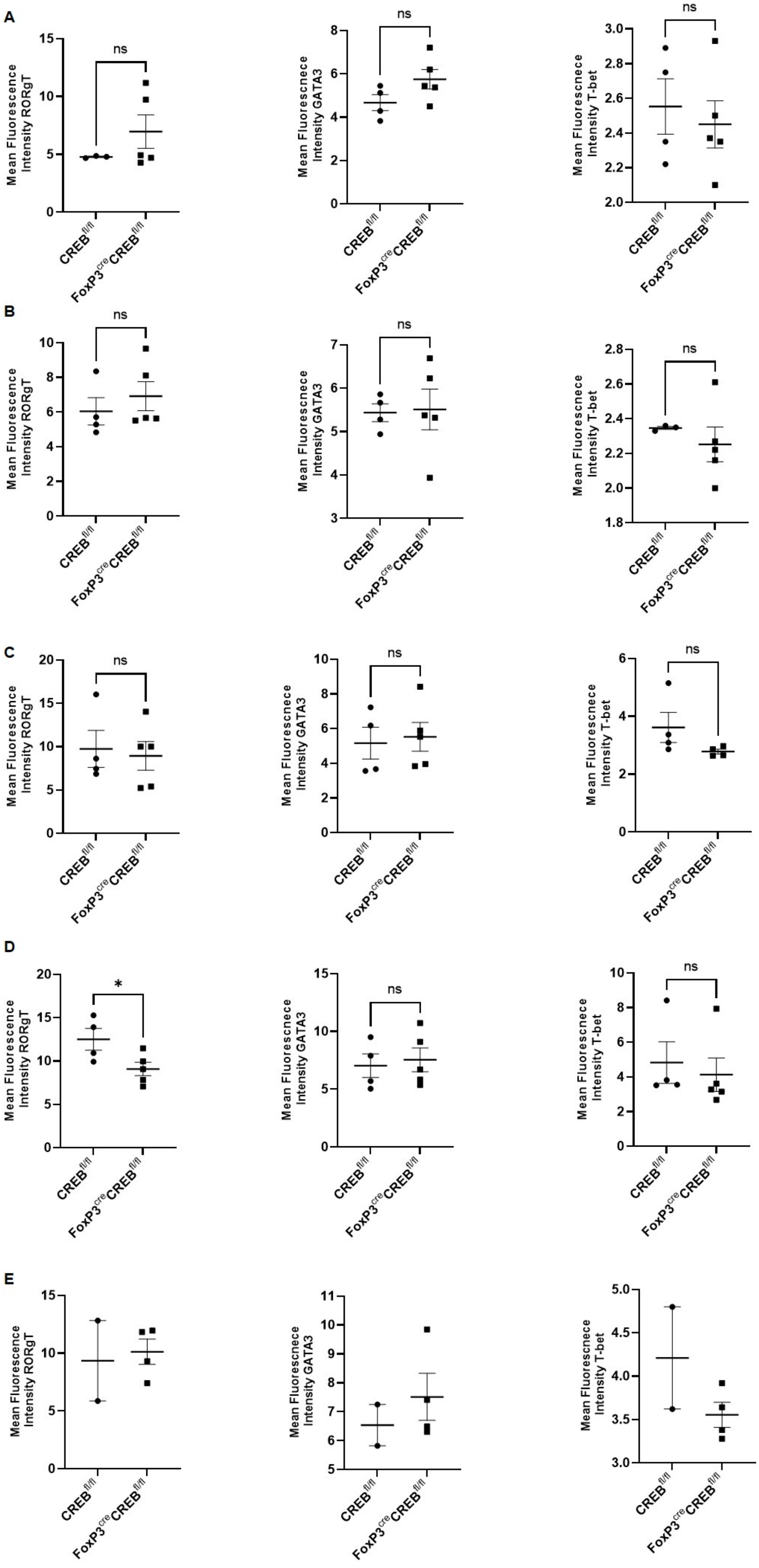
A
B
C
D
E
Mean Fluorescence Intensity RORgT
Mean Fluorescnece Intensity GATA3
Mean Fluorescnece Intensity T-bet
ns
*
CREB fl/fl
FoxP3 cre CREB fl/fl

**Supplementary Figure 2: Expression of $T_h$-lineage transcription factors in different organs.** Mean-fluorescent intensity of RORγt, GATA3 and T-bet expression in $CD45^+CD4^+Foxp3^+$ cells in **A)** Spleen, **B)** Mesenteric lymph node, **C)** Lung, **D)** Liver, and **E)** Colon. Each dot represents one animal. ns - $p>0.05$, $*p<0.05$ and results are expressed as the mean ± SEM.

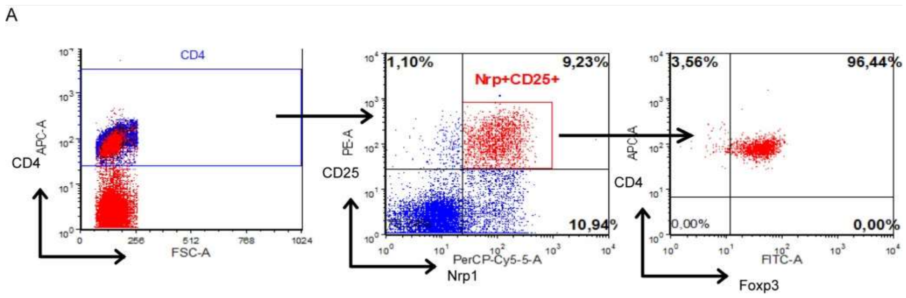


**Supplementary Figure 3: Gating strategy to sort $T_{reg}$ cells.** We sorted $Nrp1^{+}CD25^{+}CD4^{+}$ cells and thereby achieved a purity of 96% $Foxp3^{+}$ cells.

A

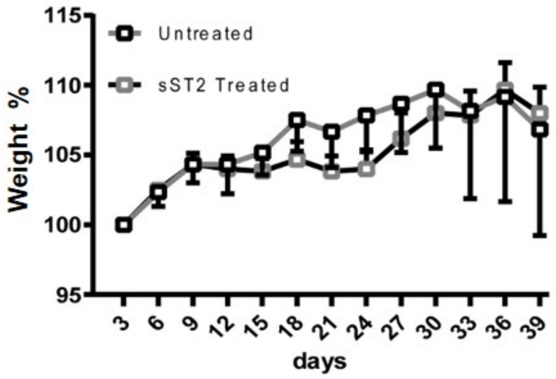


**Supplementary Figure 4: Reduced colitis in *Foxp3$^{cre}$CREB$^{fl/fl}$* CD4$^{+}$ T cell recipients is unaffected by ST2 blockade. A)** Rag2$^{-/-}$ mice were adoptively transferred with *Foxp3$^{cre}$CREB$^{fl/fl}$* CD4$^{+}$ T cells (CD4+CD25-). Mice were either treated with sST2 or PBS. Body weight is shown as percentage of starting weight (*N* = *6*, symbols indicate mean and error bars represent means ±SEM).

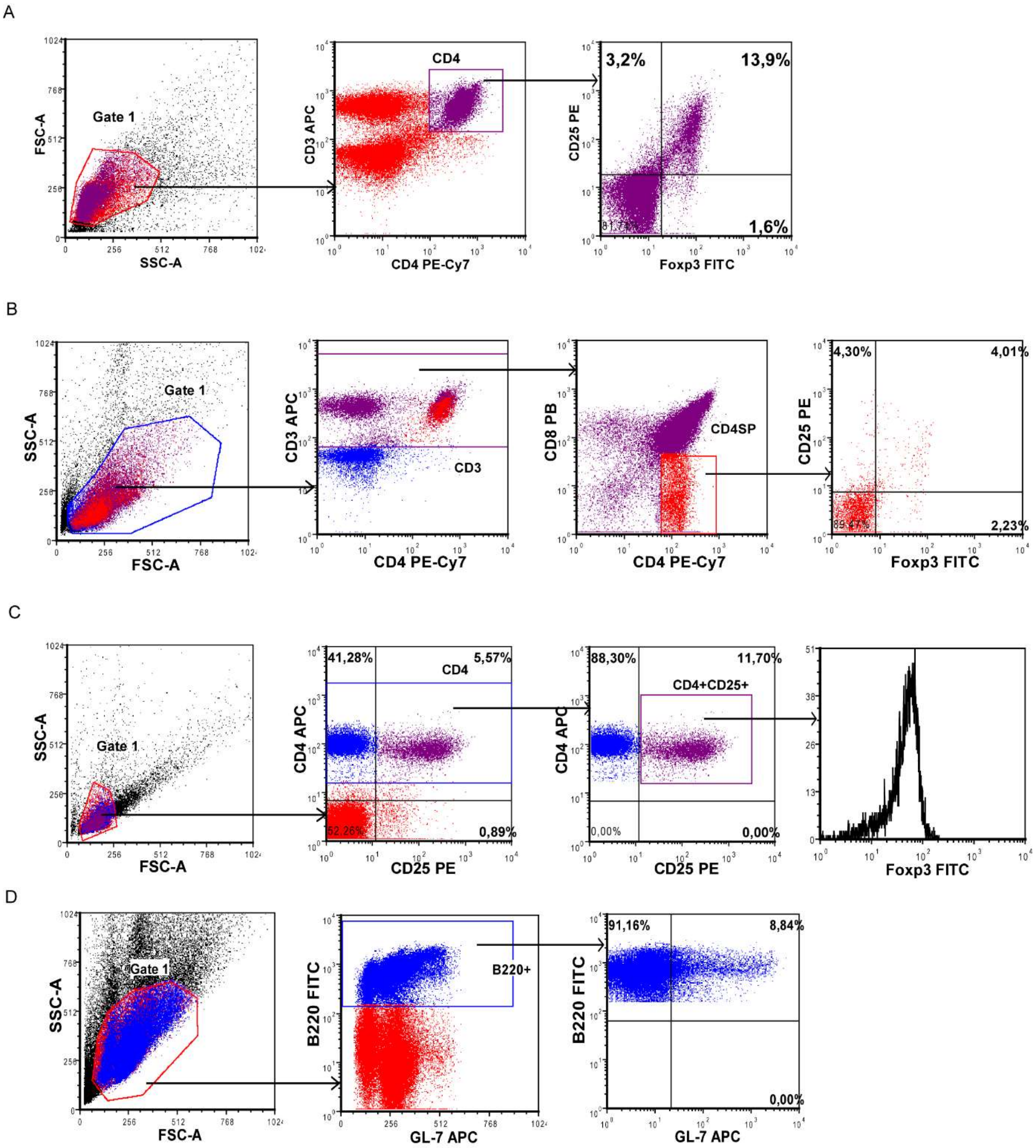


**Supplementary Figure 5: Gating strategies. A)** Figure exemplifying gating strategy to determine $CD25^+Foxp3^+$ $T_{regs}$ within LNs and spleens. **B)** Figure exemplifying gating strategy to determine $CD25^+Foxp3^+$ $T_{regs}$ within thymus. **C)** Figure exemplifying gating strategy to determine Foxp3 MFI within $CD4^+CD25^+$ cells. **D)** Figure exemplifying gating strategy to determine percentages of GL-$7^+$ cells within $B220^+$ cells.

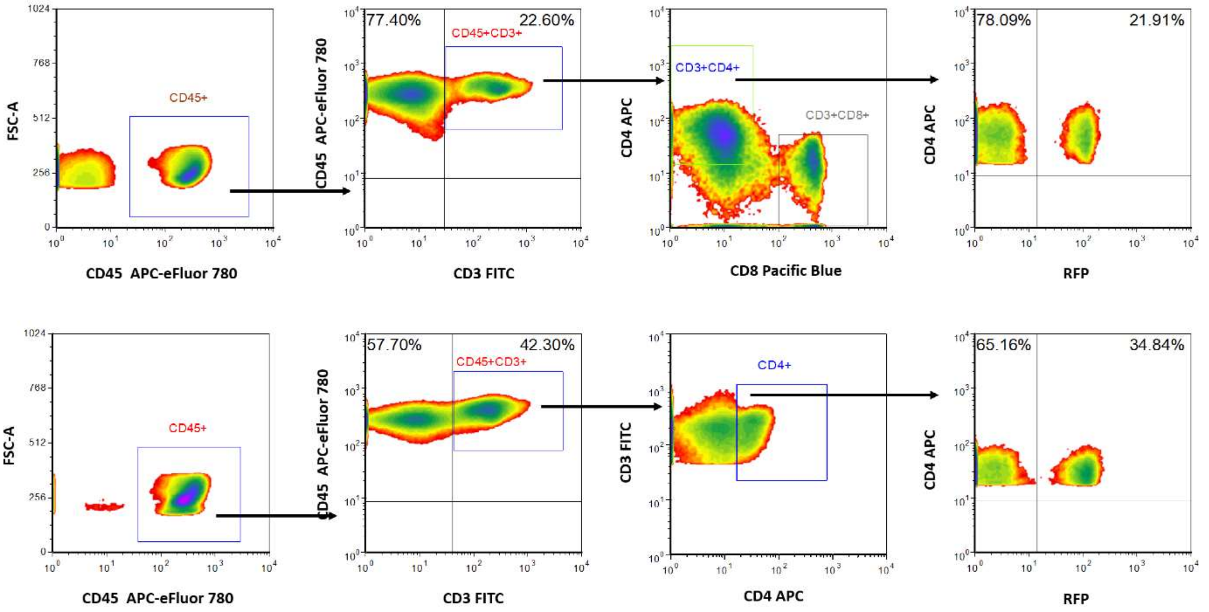


**Supplementary Figure 6: Gating strategies used in** *Foxp3$^{cre}$ROSA$^{RFP}$* mice. Figure exemplifying gating strategy to determine $T_{regs}$ within **A)** lung and **B)** colon.

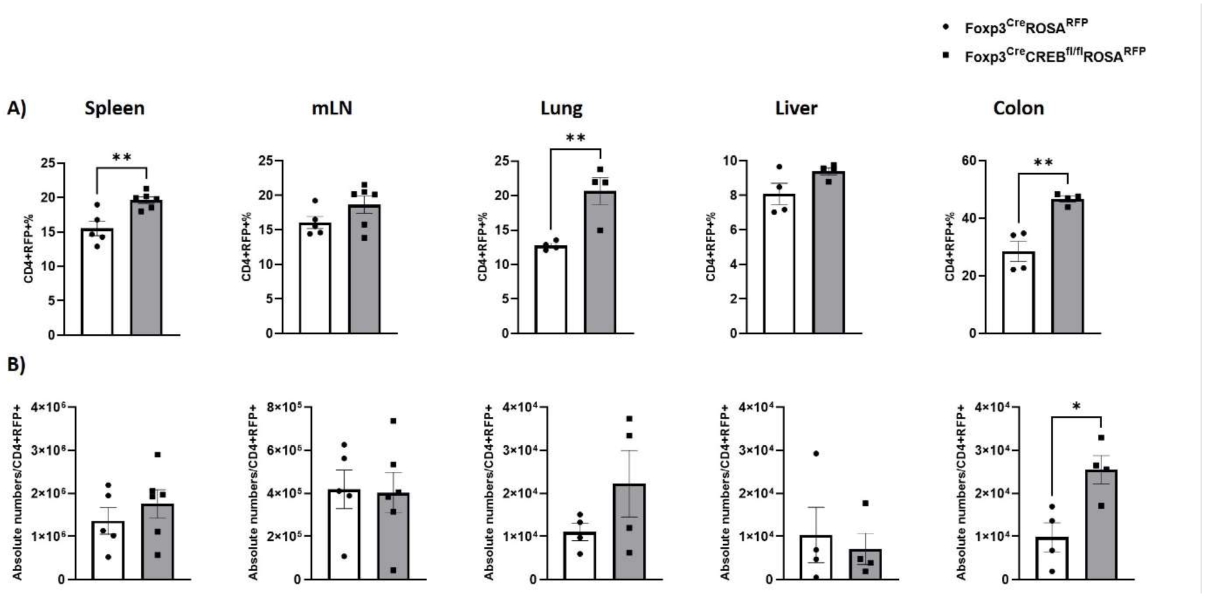


**Supplementary Figure 7:** Cell frequencies **A)** and absolute cell numbers **B)** of RFP+ cells in spleens, mLN (mesenteric lymph nodes), lungs, livers and colons of Foxp*3*$^{cre}$*ROSA*$^{RFP}$ and *Foxp3*$^{cre}$*CREB*$^{fl/fl}$*ROSA*$^{RFP}$ mice. Unpaired Student t-test was performed. **p<0.01 and results are expressed as the mean ± SEM.

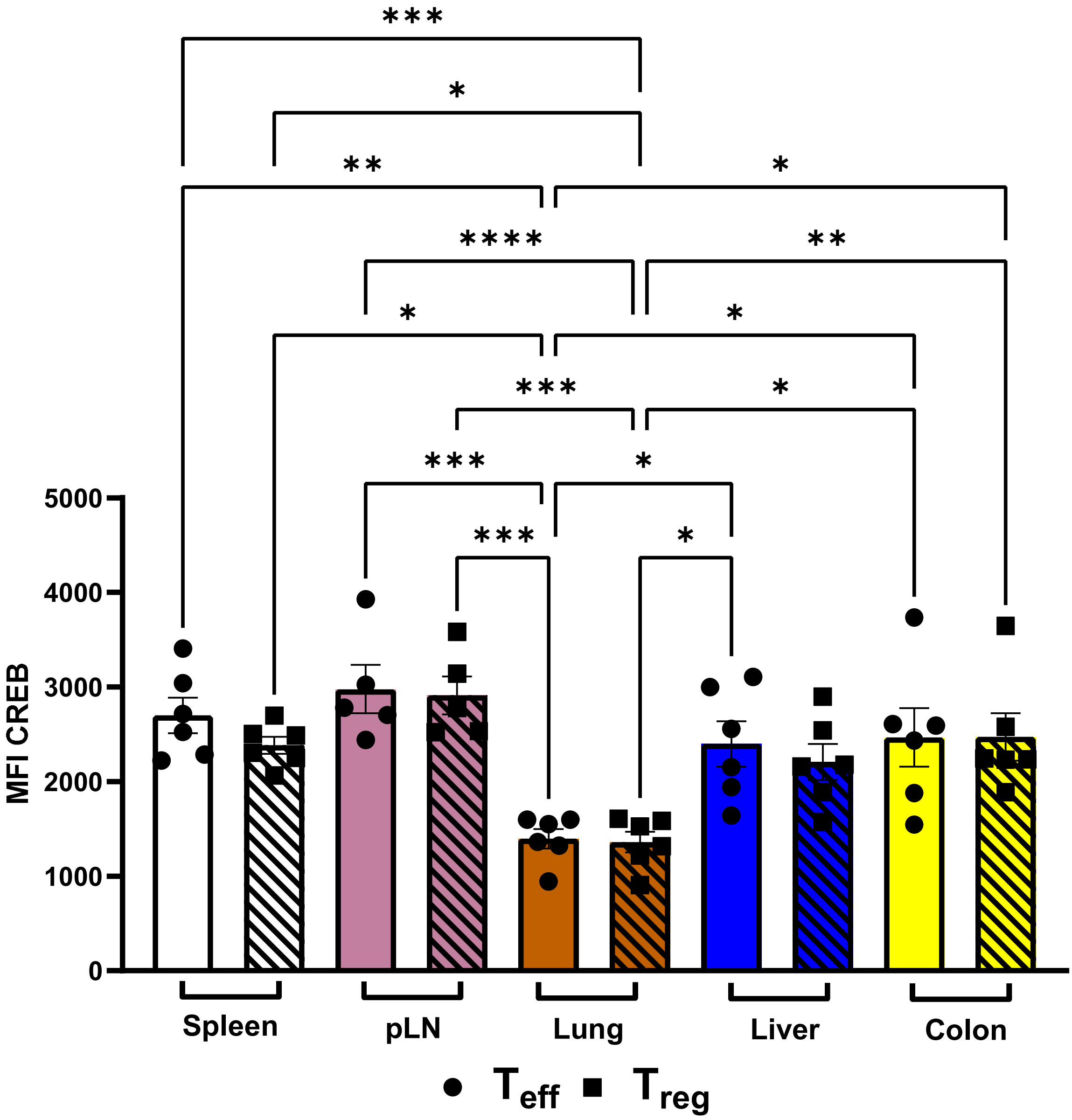


**Supplementary Figure 8:** Expression of CREB on $T_{eff}$ and $T_{reg}$ cell populations in Spleens, peripheral Lymph nodes, Lungs, Livers and colons of Foxp*3$^{cre}$ROSA$^{RFP}$* mice. ONE-way ANOVA test was performed. *p<0.05, **p<0.01, ***p<0.001, ****p<0.0001 and results are expressed as the mean ± SEM.

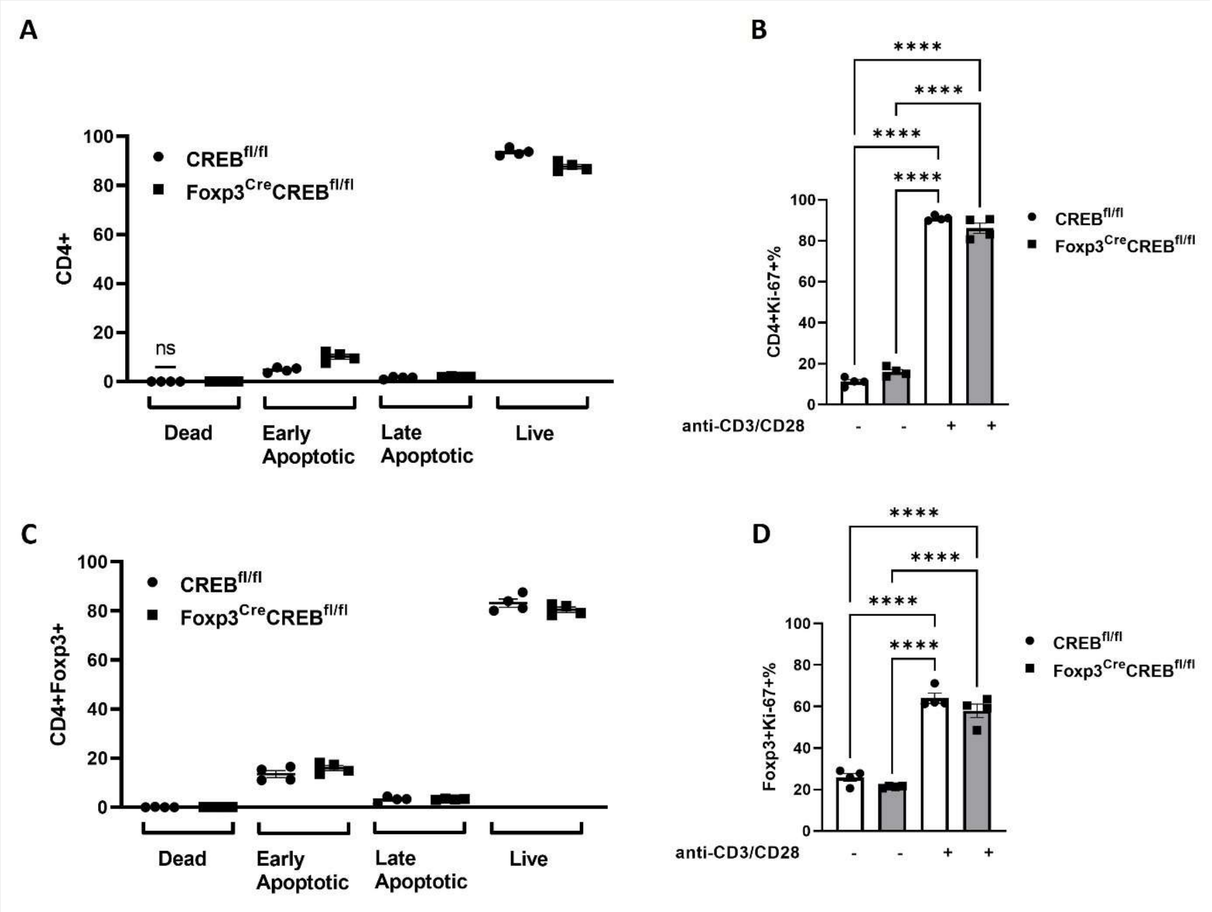


**Supplementary Figure 9:** Splenocytes of *WT and Foxp3$^{cre}$CREB$^{fl/fl}$* mice were stimulated with anti-CD3/CD28 for 2 days and stained for cell survival and proliferation. Annexin and Fixable viability-stained cells of **A)** CD4+ & **C)** CD4+Foxp3+ cells. Ki-67-stained cell percentages of **B)** CD4+ & **D)** CD4+Foxp3+ cells. ONE way-ANOVA test was performed. ns – $p>0.05$, ****$p<0.0001$ and results are expressed as the mean ± SEM.
